# TRPM2 couples cell-autonomous type-I interferon signaling to pigmentation homeostasis

**DOI:** 10.64898/2026.03.09.710699

**Authors:** Nutan Sharma, Abhishek Tanwar, Changyu Zheng, Indu Ambudkar, Rajender K Motiani

## Abstract

Transient Receptor Potential Melastatin 2 (TRPM2), a Ca²⁺-permeable cation channel, regulates innate and adaptive immunity and has recently been implicated in vitiligo, an autoimmune pigmentary disorder. However, whether TRPM2 exerts cell-autonomous immunoregulatory functions and how such signaling intersects with pigmentation remain unknown. Here, we reveal an unexpected role for TRPM2 as an intrinsic suppressor of pigmentation through type-I interferon (IFN) signaling in melanocytes. Pharmacological inhibition, genetic silencing, and gain-of-function approaches demonstrate that TRPM2 negatively regulates melanogenesis *in vitro*. Notably, TRPM2-deficient zebrafish and TRPM2⁻/⁻ mice exhibit enhanced pigmentation *in vivo*, establishing physiological relevance. Transcriptomic profiling uncovers autonomous activation of the type-I-IFN pathway upon TRPM2 loss, leading to induction of interferon-stimulated gene 15 (ISG15). Mechanistically, ISG15 attenuates global ubiquitination and stabilizes microphthalmia-associated transcription factor (MITF), the master regulator of melanogenesis, thereby promoting pigmentation. Collectively, our findings define a previously unrecognized TRPM2–type-I-IFN–ISG15–MITF signaling axis that functionally integrates cell-autonomous immune surveillance pathways with pigmentary control. Further, it provides a conceptual framework linking type-I interferon signaling to pigmentation homeostasis and pigmentary disorders.

**Key highlights of the study:**

- TRPM2 negatively regulates pigmentation *in vitro* (mouse and primary human cells) and *in vivo* (zebrafish and mice).
- Unbiased RNA-sequencing identifies ISG15 as a positive regulator of melanogenesis downstream of TRPM2 silencing.
- TRPM2 knockdown generates a cell autonomous type-I-IFN response in melanocytes that induces ISG15 expression.
- ISG15 antagonizes global ubiquitination and regulates stability of MITF, the master regulator of pigmentation.

## Introduction

Skin pigmentation is a natural process that provides protection against harmful UV rays (Motiani et al., 2018; Tanwar et al., 2024) and its disruption results in pigmentation deficits and increased susceptibility towards skin disorders and cancers (Natarajan, Ganju, Ramkumar, et al., 2014; Q. Wang et al., 2023). Pigmentation is characterized by melanin synthesis inside a specialized subcellular lysosome-derived organelle (LRO) known as melanosome (Le et al., 2021; Lin & Fisher, 2007). Melanosome biogenesis is divided into 4 stages, from immature or primary melanosomes (stage I and stage II) to fully melanized melanosomes (stage III and stage IV) (Le et al., 2021; Lin & Fisher, 2007). The process of melanogenesis is regulated by various extrinsic, intrinsic factors and signaling pathways that collectively modulate melanocyte functioning (Fu et al., 2020; Sharma et al., 2023; Vangeel & Voets, 2019; Wu et al., 2023). Among these, Ca^2+^ signaling has recently emerged as critical regulator of melanocyte biology and skin pigmentation (Ahuja et al., 2025). Intracellular Ca^2+^ acts as a ubiquitous secondary messenger that regulate melanocyte differentiation, melanosome transport, tyrosinase enzymatic activity and cellular response to external stimuli such as UV radiation (Belote & Simon, 2019; Devi et al., 2009; Joshi et al., 2007; Singh et al., 2017).

Several ion channels maintain Ca^2+^ homeostasis within melanocytes, among which Transient Receptor Potential (TRP) channels, a superfamily of non-selective voltage gated cation channels are critical mediators of intracellular Ca^2+^ dynamics in response to various stimuli (Clapham, 2003; Vangeel & Voets, 2019). Within the TRP superfamily, Transient Receptor Potential Melastatin 1 (TRPM1) has been reported to modulate pigmentation (Oancea et al., 2009). Despite growing recognition of TRP channels in physiology, the role of other TRPM family members in pigmentation biology remains poorly understood. TRPM2, belonging to the same TRPM family, serves as a redox sensor and is potently activated by oxidative stress via intracellular generation of ADP-ribose (ADPR) (Huang et al., 2018; Sumoza-Toledo & Penner, 2011; Takahashi et al., 2011). Recent studies have implicated that TRPM2 contributes to the vitiligo progression, a chronic autoimmune pigmentary disorder (P. Kang et al., 2018, 2024). These studies have demonstrated TRPM2 promotes immune cell recruitment for melanocyte destruction. While these studies underscore the pathological importance of TRPM2 in vitiligo progression, they largely focus on oxidative stress mediated cytotoxicity and it remains elusive whether TRPM2 can directly regulate the primary physiological function of melanocytes i.e. melanogenesis. Additionally, these studies majorly focused on oxidative stress and type-II IFN axis, however the role of type-I IFN in pigmentation biology remains largely unappreciated. Further contribution of any ion channel mediated-IFN signaling in melanocytes biology remains unknown.

TRPM2 has emerged as a central player of immunity and inflammation in immune cells (Beceiro et al., 2017; Knowles et al., 2011; Syed Mortadza et al., 2015; L. Wang et al., 2020; Yamamoto et al., 2008; Zong et al., 2022). Interestingly, melanocytes primarily recognized for melanin synthesis, contribute to both innate and adaptive immunity (Faught & Schaaf, 2025; Gasque & Jaffar-Bandjee, 2015; Speeckaert et al., 2022). On the other hand, studies have demonstrated that immunological molecules can regulate melanogenesis (Choi et al., 2013; Fu et al., 2020; Natarajan, Ganju, Singh, et al., 2014; Rashighi et al., 2014; Speeckaert et al., 2022). Moreover, several pigmentary disorders are also reported with immune dysfunction (Rausch et al., 2025; Tajik et al., 2024). Given the established immunomodulatory function of TRPM2 in immune cells and recognition of melanocytes as active immune contributors, it is intriguing to explore whether TRPM2 can regulate immune surveillance pathways in melanocytes and how TRPM2 dependent immunomodulation influences melanogenesis.

Here, we reveal a novel immunomodulatory function of TRPM2 in melanocytes, which negatively regulates melanogenesis. We utilized 2 independent *in vitro* model systems (B16 mouse melanoma cells and primary human melanocytes) to report that TRPM2 negatively regulates pigmentation. Further, we used 2 independent *in vivo* model systems (TRPM2 zebrafish morphants and TRPM2^-/-^ mice) to validate our *in vitro* findings. By utilizing pharmacological inhibition, siRNA-mediated TRPM2 silencing and TRPM2 overexpression, we reveal that TRPM2 negatively regulates pigmentation *in vitro*. Further, TRPM2 knockdown in zebrafish larvae showed increased pigmentary phenotype. Moreover, TRPM2^-/-^ mice showed a significant increase in epidermal tail pigmentation as compared to wild-type (WT) mice. Collectively, this data establishes TRPM2 as a critical negative regulator of pigmentation. To understand the downstream molecular mechanism, we performed unbiased RNA-sequencing. We identified that TRPM2 silencing activates type-I-IFN response that subsequently activates IFN stimulated gene 15 (ISG15). ISG15 knockdown and overexpression studies validate its crucial role in pigmentation downstream of TRPM2. Mechanistically, we show that ISG15 positively regulates MITF expression and that in turn drives pigmentation. We demonstrate that ISG15 inhibits ubiquitination, which subsequently enhance stability of MITF and other melanogenic proteins. Taken together, our study unveils a critical immunomodulatory function of TRPM2 in melanocytes wherein its silencing generates a cell autonomous type-I-IFN response. This in turn activates ISG15, which regulates MITF expression to govern the process of melanogenesis.

## RESULTS

### TRPM2 negatively regulates pigmentation *in vitro*

We first investigated TRPM2 expression during pigmentation induction in the well-established B16 mouse cell line based low-density (LD) pigmentation model (Motiani et al., 2018; Natarajan, Ganju, Singh, et al., 2014). In this model, low-density plating of depigmented B16 cells leads to autonomous and synchronous pigmentation over a period of 6-7 days (**Fig. 1A**).

**Fig 1:**
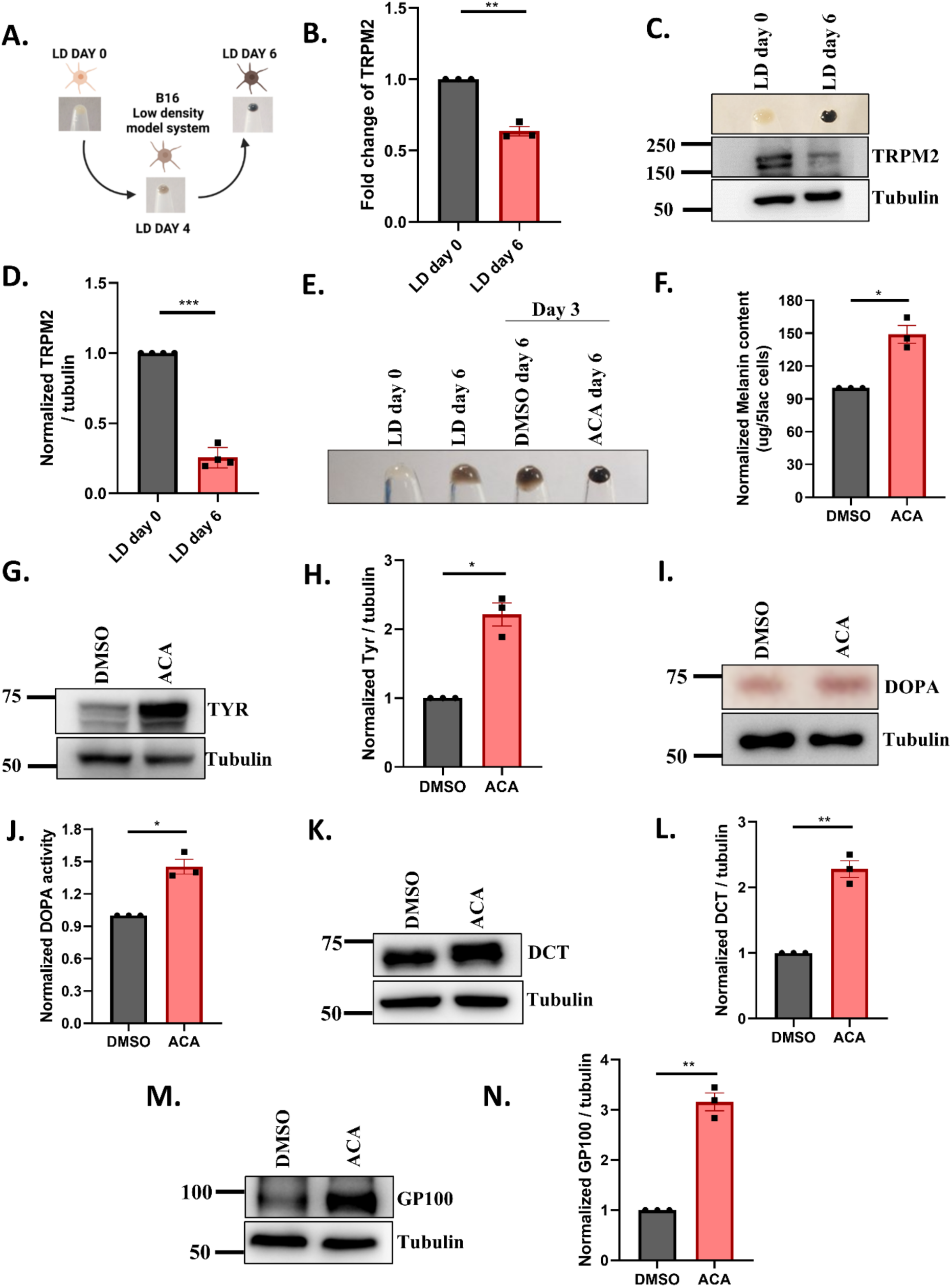
TRPM2 expression is negatively associated with pigmentation. (A) Representative B16 cell pellet images from LD day 0, LD day 4 and LD day 6 (*N* = 3) (B) qRT-PCR analysis showing TRPM2 mRNA levels in LD day 0 and LD day 6 cells (N = 3). (C) Representative western blot showing TRPM2 protein levels in LD day 0 and LD day 6 cells (N=4). (D) Densitometric quantitation of TRPM2 protein levels in LD day 0 and LD day 6 cells (N=4). (E) Representative B16 pellet images showing (Left to right) LD day 0, WT LD day 6, DMSO treated LD day 6, ACA treated LD day 6 cells (N=3). (F) Quantitation of melanin content from DMSO and ACA treated LD day 6 cells (N=3). (G) Representative western blot of TYR from DMSO and ACA treated LD day 6 cells (N=3). (H) Densitometric quantitation of TYR protein levels from DMSO and ACA treated LD day 6 cells (N=3). (I) Representative in-gel image of DOPA assay from DMSO and ACA treated LD day 6 cells (N=3). (J) Densitometric quantitation of DOPA assay from DMSO and ACA treated LD day 6 cells (N=3). (K) Representative western blot of DCT from DMSO and ACA treated LD day 6 cells (N=3). (L) Densitometric quantitation of DCT protein levels from DMSO and ACA treated LD day 6 cells (N=3). (M) Representative western blot of GP100 from DMSO and ACA treated LD day 6 cells (N=3). (N) Densitometric quantitation of GP100 protein levels from DMSO and ACA treated LD day 6 cells (N=3).

In this model, we observe moderate pigmentation at day 4 and hyperpigmentation by day 6. To evaluate the association of TRPM2 with pigmentation, we first determined the changes in TRPM2 mRNA levels by performing qRT-PCR in B16 LD model. We observed a significant reduction in TRPM2 mRNA levels on day 6 compared to day 0, suggesting an inverse relation between TRPM2 expression and melanogenesis (**Fig. 1B**). We further examined the TRPM2 protein expression in these cells. Interestingly, we observed a significant reduction in TRPM2 protein level on LD day 6 compared to day 0 (**Fig. 1C-D**). This data demonstrates that TRPM2 is inversely associated with pigmentation.

We next started studying the role of TRPM2 in pigmentation. We first used N-(p-amylcinnamoyl) anthranilic acid (ACA), a pharmacological inhibitor of TRPM2 (Kraft et al., 2006; Zhang et al., 2021). ACA is reported to inhibit TRPM2 activity at 20µM concentration (Kraft et al., 2006; Zhang et al., 2021). Therefore, we treated B16 cells with either 20µM ACA or vehicle control DMSO on LD day 3 and harvested the cells on LD day 6. Interestingly, we observed a significant increase in the pigmentation phenotype upon pharmacological inhibition of TRPM2, as evident in LD day 6 pellet images (**Fig. 1E**). We further quantitated these phenotypic changes by performing melanin content assay. We observed that TRPM2 inhibition leads to around 50% increase in the melanin levels (**Fig. 1F**). The process of melanogenesis is regulated by three pivotal proteins namely tyrosinase (TYR), dopachrome tautomerase (DCT) or TYR-related protein 2 (TYRP2/TRP2) and PMEL (also known as PMEL17 or GP100) (Le et al., 2021). Thus, we started examining the expression of these proteins upon TRPM2 inhibition. We observed a significant increase in TYR expression upon ACA treatment compared to DMSO (**Fig. 1G-H**). TYR catalyzes the rate-limiting hydroxylation of L-tyrosine to L-3,4-dihydroxyphenylalanine (L-DOPA) and the subsequent oxidation to dopaquinone within melanosomes (Le et al., 2021). Thus, we next determined the TYR enzymatic activity by performing DOPA assay and observed that TYR activity was significantly higher upon ACA treatment compared to DMSO (**Fig. 1I-J).** We further evaluated the DCT protein expression and found significant increase in its expression upon TRPM2 inhibition (**Fig. 1K-L**). Next, we assessed changes in GP100 levels, which is an integral membrane glycoprotein providing the structural scaffold for melanin deposition within melanosomes (Le et al., 2021). We observed a significant increase in GP100 protein expression upon TRPM2 inhibition with ACA as compared to DMSO control (**Fig. 1M-N**). Taken together, our results show that TRPM2 inhibition negatively regulates pigmentation.

### Gain and loss of functions studies validate that TRPM2 negatively regulates pigmentation

To validate the pharmacological inhibition data, we performed TRPM2 loss and gain of functions studies. First of all, we performed transient knockdown of TRPM2 using TRPM2 siRNA and validated TRPM2 silencing via qRT-PCR. We found a significant reduction in TRPM2 mRNA levels suggesting a robust knockdown (**Fig. 2A**). We further validated this at the protein level and we observed a significant reduction in TRPM2 protein in siTRPM2 condition in comparison to non-targeting control siRNA (siNT) (**Fig. 2B-C**). Next, we transfected B16 cells with either TRPM2 siRNA or siNT control on LD day 3 and assessed the pigmentation levels on LD day 6. As expected, we observed a robust increase in the pigmentation phenotype upon transient silencing of TRPM2 as compared to siNT control (**Fig. 2D**). We further quantitated the change in pigmentation by measuring melanin content and observed around 50% increase in melanogenesis (**Fig. 2E**). To study the changes at the molecular level, we next examined TYR expression and its activity. We found that TYR expression and activity were significantly higher upon TRPM2 silencing as compared to siNT (**Supplementary Fig. 1A-D**). We next examined changes in the DCT and GP100 protein levels and observed a significant increase in their expression upon TRPM2 silencing as compared to siNT (**Supplementary Fig. 1E-H**). To validate B16 data, we next performed TRPM2 loss of function studies in primary human melanocytes. We observed that TRPM2 silencing in primary melanocytes led to an increase in pigmentation phenotype corroborating its negative control over pigmentation (**Fig. 2F-G**). Taken together, our findings from TRPM2 silencing in B16 LD model system and primary melanocytes mirrored our observations with the TRPM2 inhibition thereby substantiating that TRPM2 serves as a negative regulator of pigmentation.

**Fig 2:**
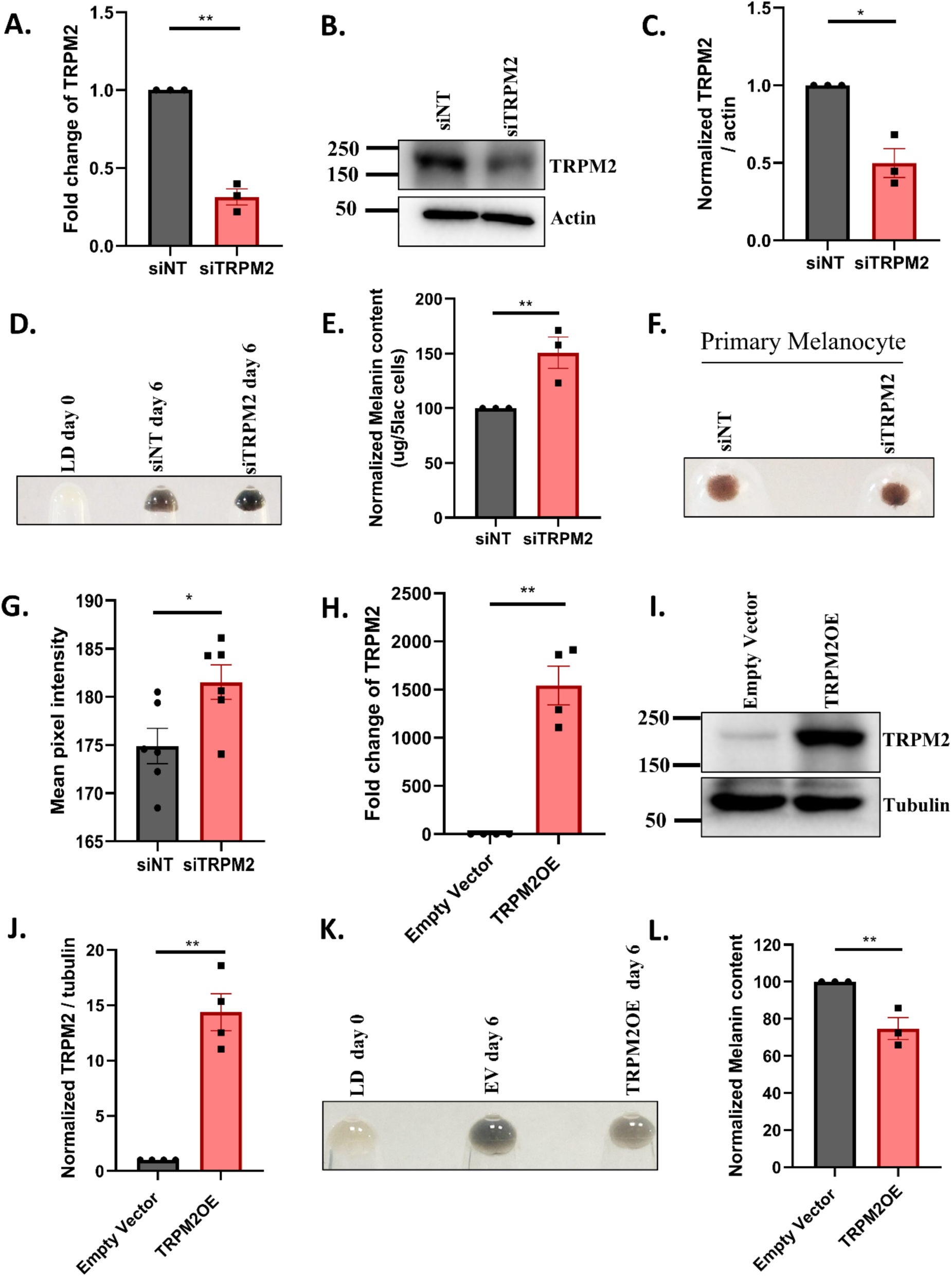
TRPM2 loss and gain of function studies establish it as a negative regulator of pigmentation. (A) qRT-PCR analysis showing TRPM2 mRNA levels from either siNT or siTRPM2 transfected B16 cells (N=3). (B) Representative western blot of TRPM2 protein levels from either siNT or siTRPM2 transfected LD day 6 cells (N=3). (C) Densitometric quantitation of TRPM2 protein levels from either siNT or siTRPM2 transfected LD day 6 cells (N=3). (D) Representative B16 pellet images from LD day 0, siNT LD day 6 and siTRPM2 LD day 6 cells (N=3). (E) Quantitation of the melanin content assay from either siNT or siTRPM2 transfected LD day 6 cells (N=3). (F) Representative pellet images from either siNT or siTRPM2 transfected primary melanocytes (N=6). (G) Mean pixel intensity quantitation of pellets images from either siNT or siTRPM2 transfected primary melanocytes (N=6) (H) qRT-PCR analysis showing TRPM2 mRNA levels in either EV control or TRPM2 transfected LD day 6 cells (N=4). (I) Representative western blot showing TRPM2 protein levels from either EV control or TRPM2 transfected LD day 6 cells (N=4). (J) Densitometric quantitation of TRPM2 protein levels from either EV control or TRPM2 transfected LD day 6 cells (N=4). (K) Representative B16 pellet images from LD day 0, either EV control or TRPM2 transfected LD day 6 cells (N=3). (L) Quantitation of the melanin content assay from either EV control or TRPM2 transfected LD day 6 cells (N=3).

To further confirm TRPM2’s role in pigmentation, we conducted gain-of-function experiments. We first performed TRPM2 overexpression (OE) using TRPM2 plasmid and validated the OE at both mRNA and protein levels. We observed a significant increase in the TRPM2 mRNA levels upon OE as compared to empty vector (EV) control (**Fig. 2H**). Similarly, we also observed a significant increase in TRPM2 protein levels (**Fig. 2I-J**). We then transfected B16 cells with either TRPM2 plasmid or with the EV control on LD day 3 and harvested cells at LD day 6. Excitingly, we observed a substantial decrease in the pigmentation phenotype following TRPM2 OE as compared to EV control (**Fig. 2K**). To further validate this phenotype, we quantitated the melanin and found a significant reduction in melanin content upon TRPM2 OE as compared to EV control (**Fig. 2L**). We next investigated whether the reduction in pigmentation upon TRPM2 OE was associated with changes in key melanogenic protein levels. We thus examined the expression of TYR, DCT and GP100 upon TRPM2 OE by performing western blotting. As expected, we observed that TRPM2 OE leads to a decrease in the TYR expression and its enzymatic activity (**Supplementary Fig. 2A-D**). Further, we also observed significant reduction in DCT and GP100 protein levels upon TRPM2 OE. (**Supplementary Fig. 2E-H**). Collectively, our data with TRPM2 pharmacological inhibition, TRPM2 gain & loss of function studies in B16 LD pigmentation model and TRPM2 silencing in primary human melanocytes elegantly demonstrate that TRPM2 is a negative regulator of the pigmentation. Further, downstream of TRPM2 silencing, increase in the pigmentation phenotype is directly associated with augmented expression of key melanogenic proteins.

### TRPM2 negatively regulates pigmentation *in vivo*

We next performed *in vivo* experiments to validate TRPM2’s role in regulating pigmentation wherein we first utilized zebrafish model system. We silenced TRPM2 in zebrafish using morpholinos specifically targeting TRPM2 and used a scrambled morpholino as control (**Fig.** 3A) (Please refer to methods section for details of zebrafish injections). Morpholinos or MOs are short synthetic antisense oligonucleotides that are used in zebrafish to block gene expression. We performed TRPM2 silencing using TRPM2 MO and validated the knockdown with qRT-PCR. We observed a significant reduction in TRPM2 mRNA levels (**Fig. 3B**). We next observed that TRPM2 morphants showed increased pigmentary phenotype compared to control MO corroborating that TRPM2 is a negative regulator of pigmentation (**Fig. 3C**). We quantitated the differences in pigmentation phenotype using ImageJ software and observed a significant increase in mean pixel intensity upon TRPM2 silencing (**Fig. 3D**). We next performed the melanin content assay on zebrafish embryos (Saurav & Motiani, 2025; Tanwar et al., 2024) and observed a robust over 40% increase in melanin upon TRPM2 silencing (**Fig. 3E**). Moreover, we validated TRPM2’s role in pigmentation by analyzing the epidermal (tail) pigmentation in TRPM2 global knockout (TRPM2^-/-^) mice. In comparison to wild type (WT) mice, TRPM2^-/-^ mice showed augmented tail epidermal pigmentation phenotype (**Fig. 3F**). Next, we quantitated these pigmentation differences using ImageJ software and observed a significant increase in the pigmentation in TRPM2^-/-^ mice compared to WT mice (**Fig. 3G**).

**Fig 3:**
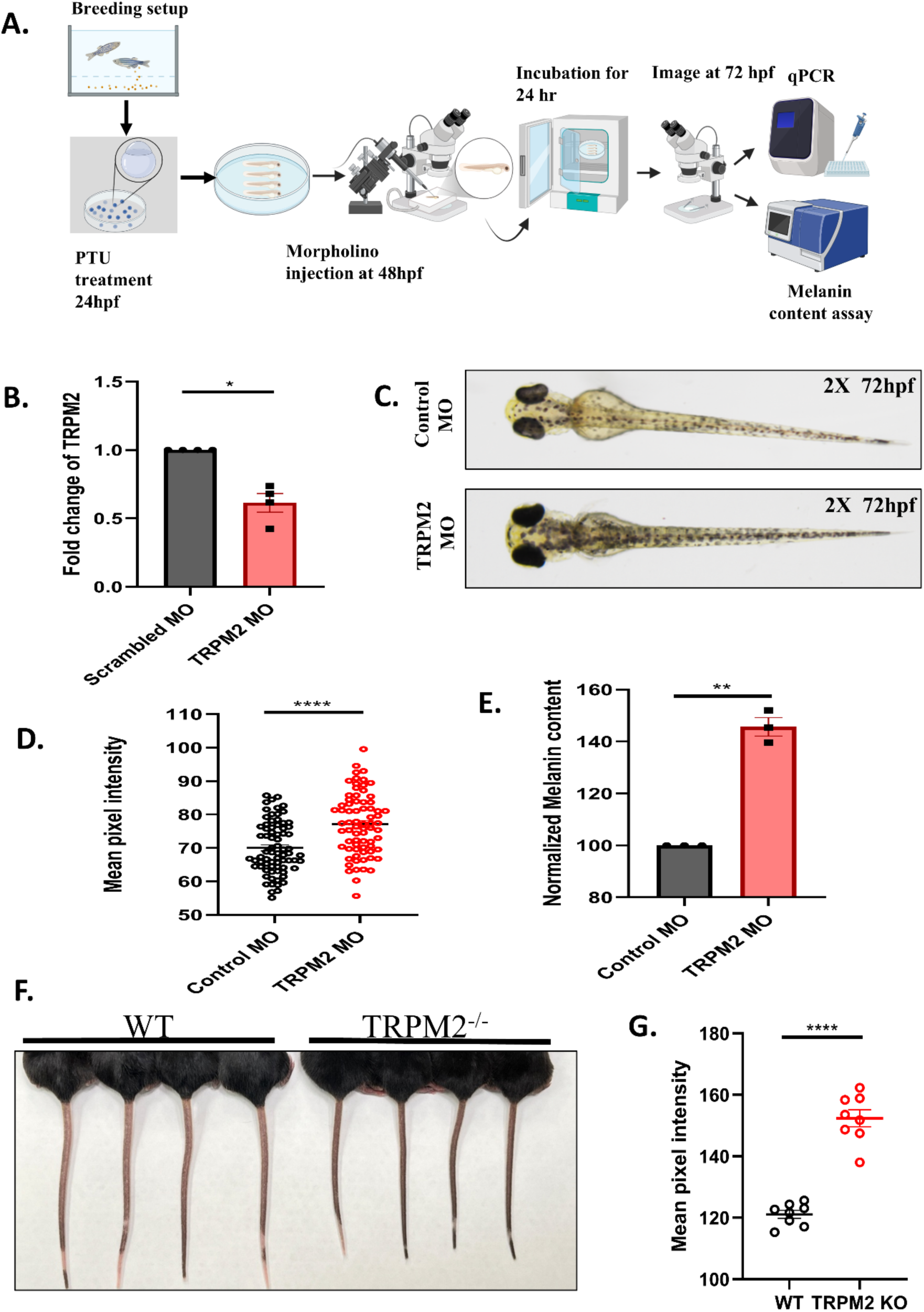
TRPM2 negatively regulates pigmentation *in vivo*. (A) Representative diagram of morpholino injections in zebrafish model system (B) qRT-PCR analysis showing TRPM2 mRNA levels in scrambled MO and TRPM2 MO conditions (N =4). (C) Representative images of zebrafish larvae at 72 hpf injected with either scrambled MO or TRPM2 MO (*N* =3 independent experiments with approximately 100 embryos/condition). (D) Quantitation of mean pixel intensity of pigmentation by ImageJ at 72 hpf in zebrafish larvae injected with either scrambled MO or TRPM2 MO (N=3 independent experiments with approximately 100 embryos/condition, each data point corresponding to one independent larvae/condition). (E) Melanin-content estimation from either scrambled MO or TRPM2 MO injected larvae (*N* =3 independent experiments with 60 embryos/condition) (F) Representative images of mice tails from approximately 3 months old WT and TRPM2^-/-^mice. (G) Quantitation of mean pixel intensity of pigmentation using ImageJ in WT mice and TRPM2^-/-^ mice (N=8, each data point corresponding to one independent mice/condition).

Collectively, our findings from 2 independent *in vitro* models (B16 cells and primary human melanocytes) and 2 distinct *in vivo* models, TRPM2 zebrafish morphants and TRPM2^-/-^ mice establish TRPM2 as a negative regulator of pigmentation.

### Unbiased RNA sequencing identifies ISG15 downstream of TRPM2

To elucidate the precise molecular mechanism regulating melanogenesis downstream of TRPM2, we conducted unbiased transcriptome profiling through RNA-seq. The sequencing was performed on TRPM2 silenced B16 LD day 6 cells and corresponding siNT control cells using the Illumina NovaSeq platform (**Fig. 4A**). We assessed the data quality utilizing 2 independent software, FastQC and MultiQC software, and subsequently aligned QC approved reads to the indexed Mus musculus genome (GRCm39) using the STAR v2 aligner. Gene expression values were acquired as read counts through feature-counts software. Subsequently, we conducted a differential expression analysis employing the DESEq2 package, and normalized counts were derived. The fold change was computed for the siTRPM2 compared to siNT. Genes exhibiting an expression of more than a log2 fold change of +1 with respect to the siNT control were categorized as upregulated genes in siTRPM2 condition. Conversely, to identify downregulated genes in siTRPM2 compared to siNT, a cut-off of a log2 fold change below -1 was used. Then, we sought to identify the biological processes modulated upon TRPM2 knockdown using pathway enrichment analysis based on the Gene Ontology (GO) database. Subsequently, the enriched pathways were plotted with their respective enrichment score using the SRPLOT web server (**Fig. 4B**). Interestingly, this robust and unbiased analysis showed cellular response to IFN beta (IFN-β) as the highest enriched pathway. The other key upregulated pathways include cell-cell adhesion, regulation of membrane potential, sodium & potassium ion transport, IFN gamma (IFN**-γ**) signaling and G-protein coupled glutamate receptor signaling pathway (**Fig. 4B**). Next, we plotted significantly altered genes in these pathways using a Heatmap and a gene volcano map using their respective expression values (**Fig. 4C-D**). Volcano plot showed significantly upregulated signatures of IFN stimulated genes (ISGs) such as ISG15, DExD/H box helicase 60 (DDx60), guanylate binding protein 7(Gbp7), Gbp3, 2’-5’ oligoadenylate synthetase 2 (Oas2), IFN-induced protein with tetratricopeptide repeats 1(Ifit1), immunity-related GTPase family M member 1 (Irgm1) upon TRPM2 silencing (**Fig. 4D**).

**Fig 4:**
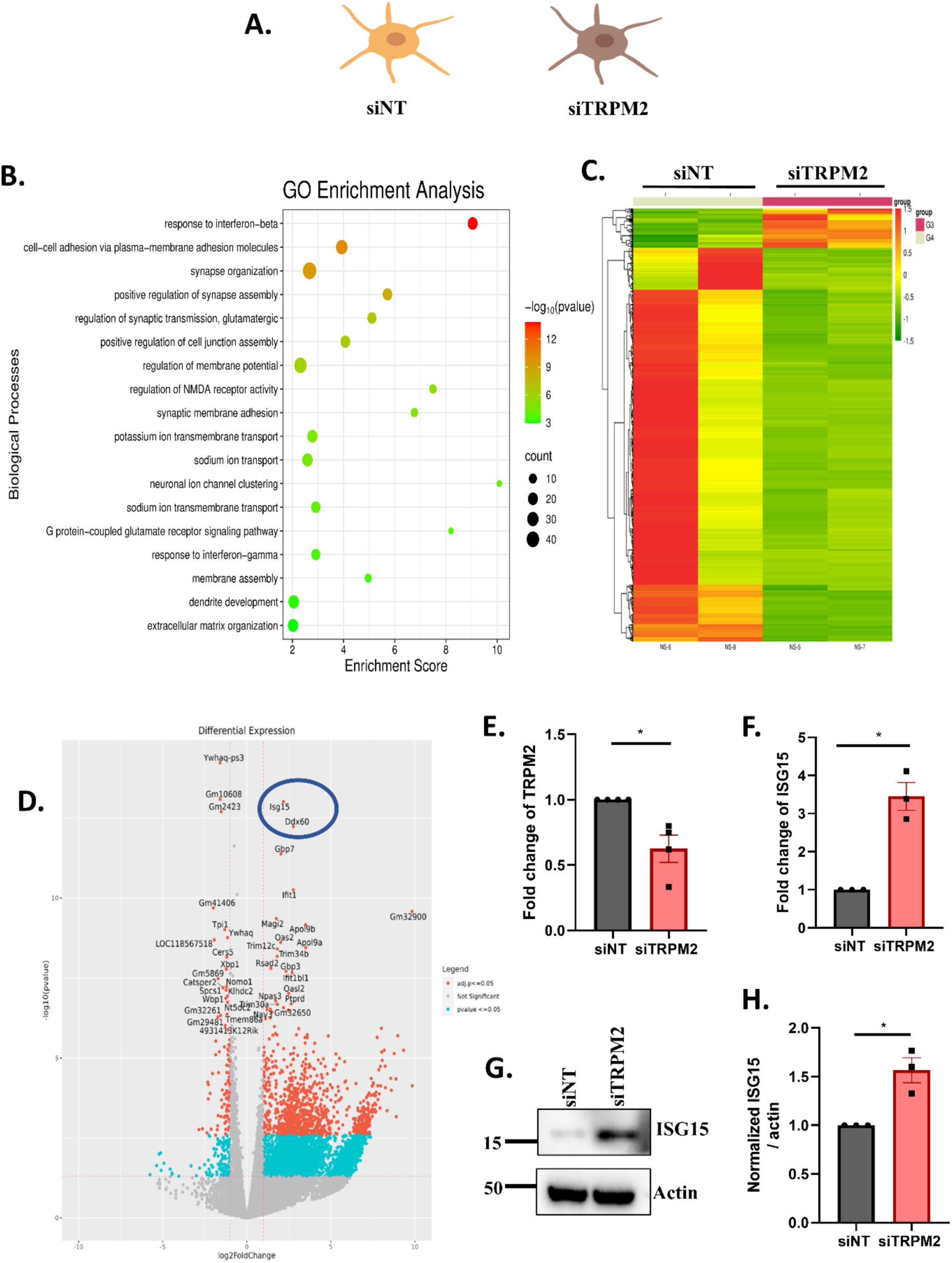
Unbiased RNA sequencing reveals ISG15 as a novel regulator of pigmentation. (A) Diagrammatic representation of siNT and siTRPM2 transfected B16 cells. (B) Gene enrichment (GO) analysis showing significant and differentially upregulated biological processes upon TRPM2 silencing. The scale on the right represents the -log_10_(p-value) from 3 to 12, labelled as green to red, respectively. (C) Heatmap representing the differential gene expression profile from either siNT or siTRPM2 transfected LD day 6 cells (N=2). The scale on the right represents the z-score for fold change from −1.5 to +1.5, labelled as green to red, respectively. (D) Volcano plot showing names of differentially regulated genes from either siNT or siTRPM2 transfected LD day 6 cells (N=2). Genes labelled in red have adjusted p-values <= 0.05, genes labelled in blue have p-values <= 0.05, and genes labelled in grey have non-significant p-values. (E) qRT-PCR analysis showing TRPM2 mRNA levels in siNT and siTRPM2 LD day 6 cells (N=4). (F) qRT-PCR analysis showing ISG15 mRNA levels in siNT and siTRPM2 LD day 6 cells (N=3). (G) Representative western blot image showing ISG15 protein levels in siNT and siTRPM2 LD day 6 cells (N=3). (H) Densitometric quantitation of ISG15 protein levels in siNT and siTRPM2 LD day 6 cells (N=3).

These upregulated ISG signatures majorly correspond to type-I-IFN pathway and innate immune response (J. A. Kang et al., 2022; McNab et al., 2015; Mirzalieva et al., 2022), which were upregulated in GO pathway enrichment analysis as well (**Fig. 4B**). This data suggests that TRPM2 silencing triggers a cell autonomous immune response, particularly type-I-IFN pathway within melanocytes. Among these immune signatures, IFN stimulated gene15 (ISG15) showed highest increase at the gene expression level. Recent studies have reported ISG15’s involvement in skin homeostasis where it is associated with intracranial calcifications and ulcerative & necrotizing skin lesions (Buda et al., 2020; Y. Liu et al., 2023; Malik et al., 2022; Martin-Fernandez et al., 2020; Oyedepo et al., 2023). However, role of ISG15 in pigmentation remains unappreciated. Therefore, we decided to investigate role of ISG15 in pigmentation.

First of all, we validated RNA-seq data by performing qRT-PCRs with the RNA-sequencing samples. We confirmed TRPM2 silencing in the sequencing samples (**Fig. 4E**). Further, we examined ISG15 mRNA levels upon TRPM2 knockdown and observed over 3-fold increase in ISG15 expression (**Fig. 4F**). Next, we analyzed ISG15 protein levels upon TRPM2 silencing and saw a significant increase in the ISG15 protein levels upon TRPM2 knockdown compared to siNT control (**Fig. 4G-H)**. This data validates that TRPM2 silencing upregulates ISG15 expression at mRNA and protein levels.

### ISG15 acts as a positive regulator of pigmentation

To delineate ISG15’s function in pigmentation, we first investigated ISG15 protein expression in B16 LD model system using LD day 0 and LD day 6 cells. We observed a significant increase in ISG15 protein levels in LD day 6 pigmented cells compared to day 0 depigmented cells (**Fig. 5A-B**). This suggested that ISG15 could be positively associated with pigmentation. Next, we conducted ISG15 silencing using siRNA and validated the knockdown at both mRNA and protein levels. We observed a substantial decrease in the ISG15 mRNA and protein levels (**Fig. 5C-E**). Subsequently, we transfected B16 cells with either ISG15 siRNA or siNT on LD day 3 and evaluated the pigmentation phenotype on LD day 6. As anticipated, we observed a significant reduction in the pigmentation phenotype in siISG15 condition compared to siNT control (**Fig. 5F**). We further assessed melanin content and observed 50% reduction in melanin upon ISG15 knockdown (**Fig. 5G**). We next assessed the changes in the melanogenic protein levels and found a significant decrease in TYR and DCT proteins in siISG15 condition compared to siNT (**Fig. 5H-K**). Thus, ISG15 silencing decreases tyrosinase and DCT levels, which in turn regulates pigmentation phenotype. We further extended our study by performing ISG15 overexpression (OE) using ISG15 plasmid. We first validated the ISG15 OE at both mRNA and protein levels using qRT-PCR and western blotting. We observed a significant increase in ISG15 expression at both mRNA and protein levels (**Fig. 5L-N**). We then transfected B16 cells with ISG15 plasmid or Empty Vector (EV) control on LD day 3 and assessed pigmentation phenotype on LD day 6. As expected, we observed a substantial increase in the pigmentation phenotype upon ISG15 OE compared to EV control (**Fig. 5O**). Further, we analyzed the pellet images from ISG15 OE & EV to assess the melanin content using ImageJ software and observed a significant increase in the mean intensity upon ISG15 OE compared to EV control (**Fig. 5P**). Next, to confirm that ISG15 works downstream of TRPM2 silencing to regulate pigmentation, we performed simultaneous silencing of TRPM2 and ISG15 in the LD pigmentation model. We first validated double knockdown of ISG15 and TRPM2 in the LD samples and then observed its outcome on the pigmentation (**Fig. 5Q-5R**). As TRPM2 depletion results in enhanced pigmentation, we examined the pigmentation status upon co-silencing of TRPM2 and ISG15. Excitingly, we observed a significant reduction in pigmentation phenotype upon double knockdown of ISG15 and TRPM2 as compared to siNT (**Fig. 5S**). We further quantified these phenotypic changes and observed a significant decrease in pigmentation upon double knockdown (**Fig. 5T**). This data demonstrates that the increase in pigmentation seen in TRPM2 depleted cells is rescued upon co-silencing of ISG15. Collectively, our findings from unbiased transcriptomics & its molecular validation, subsequent ISG15 silencing, ISG15 overexpression and co-knockdown of ISG15 with TRPM2 reveal that TRPM2 silencing induces ISG15 expression, which in turn drives pigmentation.

**Fig 5:**
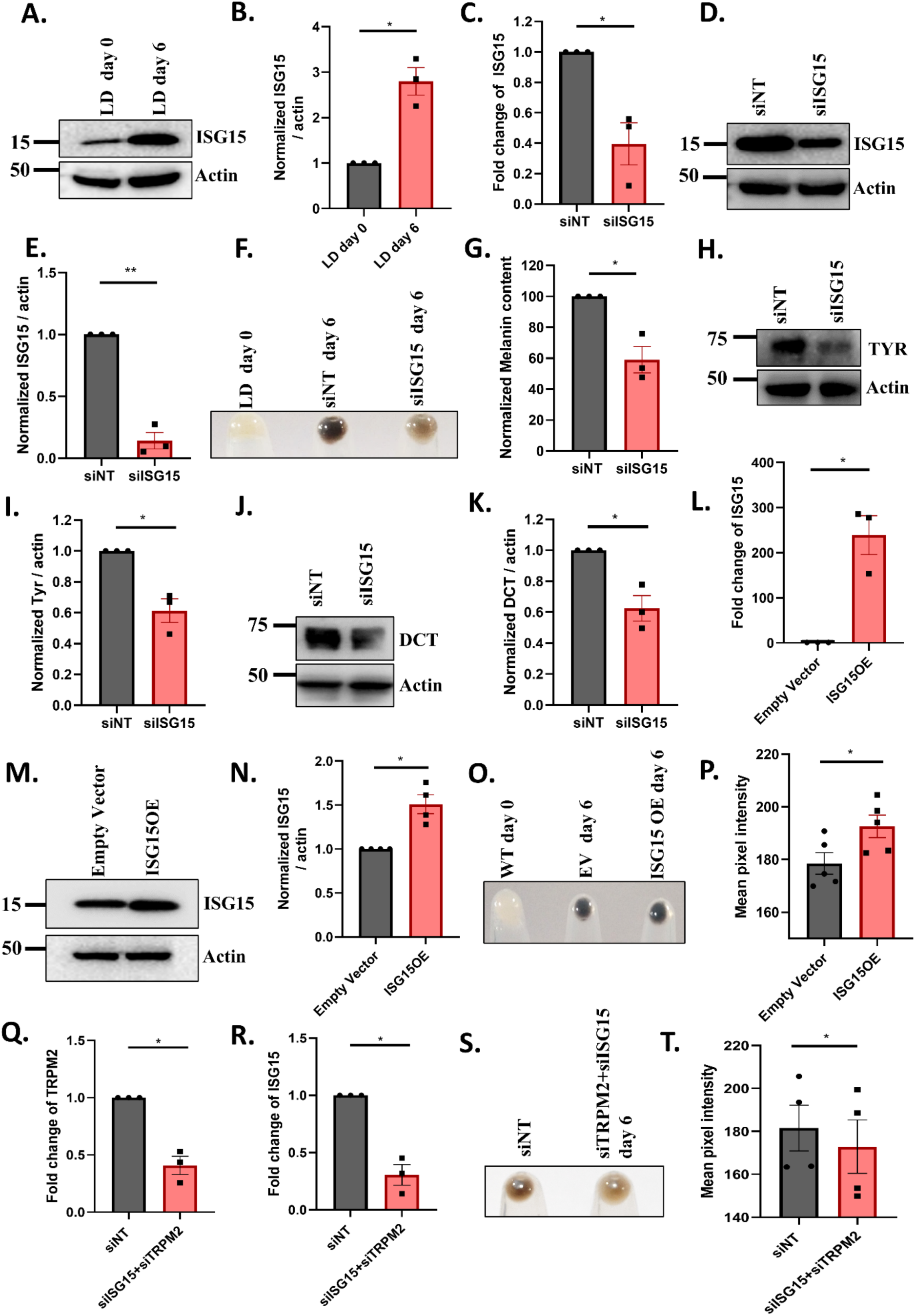
ISG15 acts as a positive regulator of pigmentation. (A) Representative western blot showing ISG15 protein levels in LD day 0 and LD day 6 cells (N=3). (B) Densitometric quantitation of ISG15 protein levels in LD day 0 and LD day 6 cells (N=3). (C) qRT-PCR analysis showing ISG15 mRNA levels in either siNT or siISG15 transfected LD day 6 cells (N=3). (D) Representative western blot showing ISG15 protein levels from either siNT or siISG15 transfected LD day 6 cells (N=3). (E) Densitometric quantitation of ISG15 protein levels in either siNT or siISG15 transfected LD day 6 cells (N=3). (F) Representative B16 pellet images of LD day 0, either siNT or siISG15 transfected LD day 6 cells (N=3). (G) Quantitation of melanin content assay from either siNT or siISG15 transfected LD day 6 cells (N=3). (H) Representative western blot of TYR from either siNT or siISG15 transfected LD day 6 cells (N=3). (I) Densitometric quantitation of TYR protein levels from either siNT or siISG15 transfected LD day 6 cells (N=3). (J) Representative western blot of DCT from either siNT or siISG15 transfected LD day 6 cells (N=3). (K) Densitometric quantitation of DCT protein levels from either siNT or siISG15 transfected LD day 6 cells (N=3). (L) qRT-PCR analysis showing ISG15 mRNA levels in either EV or ISG15 transfected LD day 6 cells (N=3). (M) Representative western blot showing ISG15 protein level in either EV or ISG15 transfected LD day 6 cells (N=4). (N) Densitometric quantitation of ISG15 protein levels in either EV or ISG15 transfected LD day 6 cells (N=4). (O) Representative B16 pellet images of LD day 0, in either EV or ISG15 transfected LD day 6 cells (N=5). (P) Mean pixel intensity quantitation of WT, in either EV or ISG15 transfected LD day 6 cells (N=5)

### ISG15 negatively regulates ubiquitination and contributes to pigmentation by modulating MITF levels

ISG15 is a ubiquitin like modifier, known for exerting post translational modification (PTM) i.e. ISGylation onto its targets (J. A. Kang et al., 2022; Mirzalieva et al., 2022). ISG15 has been reported to indirectly modulate protein degradation by competing with ubiquitin for ubiquitin-binding sites on the target proteins. Hence, ISG15 antagonize the ubiquitination of cellular proteins (Desai et al., 2006; Fan et al., 2015; Okumura et al., 2008).Thus, we investigated if ISG15 modulates ubiquitination pathway in B16 cells. To address this, we assessed the global ubiquitination status using western blotting upon ISG15 silencing and overexpression in LD pigmentation model. We observed that the global ubiquitination increased significantly upon ISG15 silencing compared to siNT control (**Fig. 6A and 6C**). Further, we checked the global ubiquitination upon ISG15 overexpression and observed that global ubiquitination is substantially decreased in ISG15 OE compared to the empty vector control (**Fig. 6B and 6D**). This data show that ISG15 negatively regulates protein ubiquitination in LD model.

**Fig 6:**
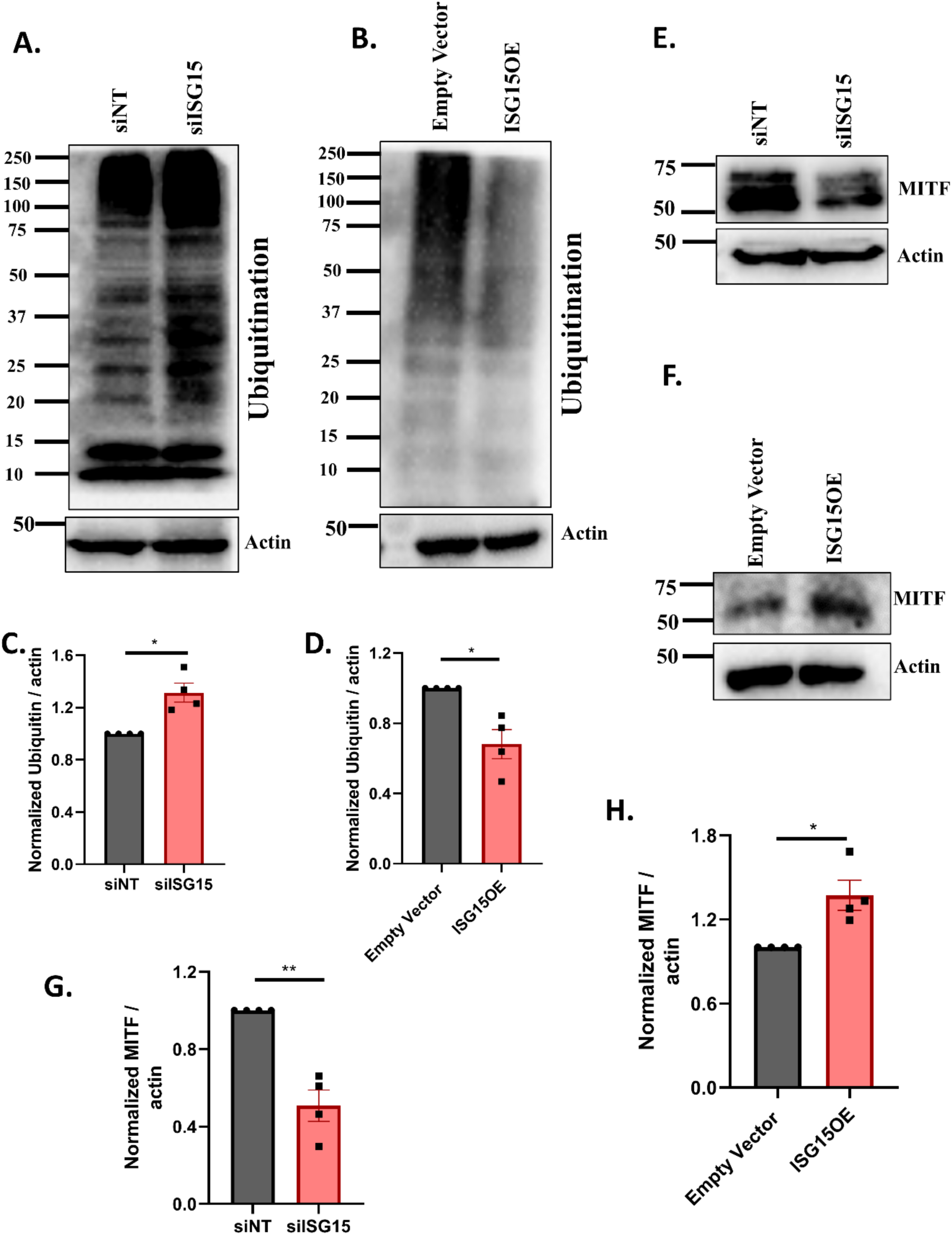
Antagonistic regulation of ubiquitination by ISG15 contributes to the pigmentation via regulating MITF degradation. (A) Representative western blot showing global ubiquitination in either siNT or siISG15 transfected LD day 6 cells (N=4). (B) Representative western blot showing global ubiquitination in either EV or ISG15 transfected LD day 6 cells (N=4). (C) Densitometric quantitation of global ubiquitination in either siNT or siISG15 transfected LD day 6 cells (N=4). (D) Densitometric quantitation of global ubiquitination in either EV or ISG15 transfected LD day 6 cells (N=4). (E) Representative western blot showing MITF protein levels in either siNT or siISG15 transfected LD day 6 cells (N=4). (F) Representative western blot showing MITF protein levels in either EV or ISG15 transfected LD day 6 cells (N=4). (G) Densitometric quantitation of MITF protein levels in either siNT or siISG15 transfected LD day 6 cells (N=4). (H) Densitometric quantitation of MITF protein levels in either EV or ISG15 transfected LD day 6 cells (N=4).

Since we observed reduction in tyrosinase and DCT expression upon ISG15 silencing (**Fig. 5H-K**), we next determined the mechanistic details of ISG15 mediated regulation of these melanogenic proteins. The expression of these proteins is driven by MITF transcription factor, the master regulator of melanogenesis. MITF is known to regulate the melanogenic genes expression, including TYR, TYRP1, MLANA, DCT and PMEL (Kawakami & Fisher, 2017). Thus, we examined MITF expression upon ISG15 silencing and observed that ISG15 silencing significantly reduces MITF protein levels compared to siNT control (**Fig. 6E and 6G**). Further, we validated ISG15’s control over MITF protein expression by overexpressing ISG15. We observed a robust increase in MITF protein levels upon ISG15 OE compared to EV control (**Fig. 6F and 6H**). Taken together, ISG15 loss and gain of function data clearly demonstrates that ISG15 positively regulates MITF expression and this altered MITF expression in turn modulates melanogenic proteins levels. Collectively, data from our unbiased RNA-seq, ISG15 silencing & OE studies demonstrate that TRPM2 silencing induces ISG15 expression and ISG15 in turn positively regulates MITF expression to augment pigmentation.

### TRPM2 silencing triggers RLR signaling and generates type-1 IFN response to activate ISG15

Since ISG15 emerged as a critical positive regulator of pigmentation downstream of TRPM2, we investigated the mechanism of ISG15 induction upon TRPM2 silencing. We analyzed the RNA-seq data to identify the genes/pathways responsible for activating the type-I-IFN response. Volcano plot data showed second highest upregulated signature of DExD/H box helicase 60 (DDx60) upon TRPM2 silencing (**Fig. 4D**). DDx60 belongs to DExD/H box helicase family and acts as key mediator of inflammation and immunity (Miyashita et al., 2011; Oshiumi et al., 2015). We first validated increase in DDx60 expression in RNA-seq samples via qRT-PCR (**Fig. 7A**). DDx60 plays a critical role in innate immunity and antiviral response by activating RLR signaling (Oshiumi et al., 2015). Interestingly, RLR signaling can induce ISG15 expression via IRF3 and IRF7 transcription factors by activating the type 1-IFN response (Lazear et al., 2019; G. Liu et al., 2021; Rehwinkel & Gack, 2020). Therefore, DDx60’s upregulation in RNA-seq led us to investigate the potential role of RLR signaling in melanogenesis. RLR belong to a class of pattern recognition receptors (PRRs), which are key sensors of both pathogens associated molecular patterns (PAMPs) and damage associated molecular patterns (DAMPs) (Li & Wu, 2021; Roh & Sohn, 2018) Recent studies have established the endogenous activation of RLR signaling in absence of any external stimuli (Chen & Hur, 2022). This pathway comprises three distinct cytoplasmic sensors namely RIG-I (encoded by gene DDx58), melanoma differentiation-associated gene 5 (MDA5) (encoded by gene IFIH1) and laboratory of genetics and physiology 2 (LGP2). These sensors are broadly expressed in a variety of cells with predominant expression in epithelial cells. We examined the RIG-I and MDA5 expression and observed increase in their mRNA levels upon TRPM2 silencing (**Fig. 7B-C**). Upon activation, RIG-I and MDA5 interact with their central downstream adaptor molecule, mitochondrial antiviral-signaling protein (MAVS) (Rehwinkel & Gack, 2020). Thus, we next examined MAVS and found upregulation in its mRNA levels upon TRPM2 silencing suggesting activation of this signaling cascade (**Fig. 7D**). MAVS activation further recruits and activates signaling molecules such as TANK-binding kinase 1 (TBK1), which in turn activates transcription factors like IFN regulatory factor IRF3 and IRF7 (Rehwinkel & Gack, 2020) thereby generating type-I-IFN response. Thus, we first assessed the TBK-1 activation upon TRPM2 silencing. We observed an increase in TBK-1 phosphorylated form while total TBK-1 protein remained unchanged suggesting its activation (**Fig. 7E-F**). Next, we followed this signaling cascade and examined the IRF3 and IRF7 activation. We observed IRF7 upregulation at both mRNA and protein levels (**Supplementary Fig. 3A and Fig. 7G)**. We further observed an increase in IRF7 phosphorylated form upon TRPM2 silencing (**Fig. 7G, Supplementary Fig. 3B**). This data showed IRF7 activation downstream of TRPM2 silencing. We next examined IRF3 which showed unchanged mRNA and endogenous protein levels (**Supplementary Fig. 3C and Fig. 7H**) however phosphorylated IRF3 was upregulated suggesting its activation (**Fig. 7H and Supplementary Fig. 3D**). Since activated IRF3 and IRF7 generate type-I-IFN response, we next estimated IFN-α and IFN-β via qRT-PCR and observed increased mRNA levels upon TRPM2 silencing (**Fig. 7I-J**). The activated IRF3 and IRF7 induce transcriptional activation of several ISGs including ISG15 to generate inflammatory and immune response (J. A. Kang et al., 2022; Mirzalieva et al., 2022). Further, to confirm the activation of type-I-IFN response, we also examined Signal Transducer and Activator of Transcription 1(STAT-1), a transcription factor and a key mediator of IFN pathway responsible for the activation of ISGs. We observed a significant increase in STAT-1 mRNA and endogenous protein levels (**Fig. 7K-L**). Furthermore, we observed an increase in STAT-1 phosphorylated form indicating type-I-IFN pathway activation (**Fig. 7L)**. Thus, our data collectively suggested that TRPM2 silencing triggers cell autonomous activation of RLR signaling cascade and thereby it generates type-I-IFN response via IRF3 and IRF7 activation within melanocytes. Subsequently, the activated IRF3 and IRF7 induce ISG15 expression, which positively regulates MITF expression to enhance melanogenesis.

**Fig 7:**
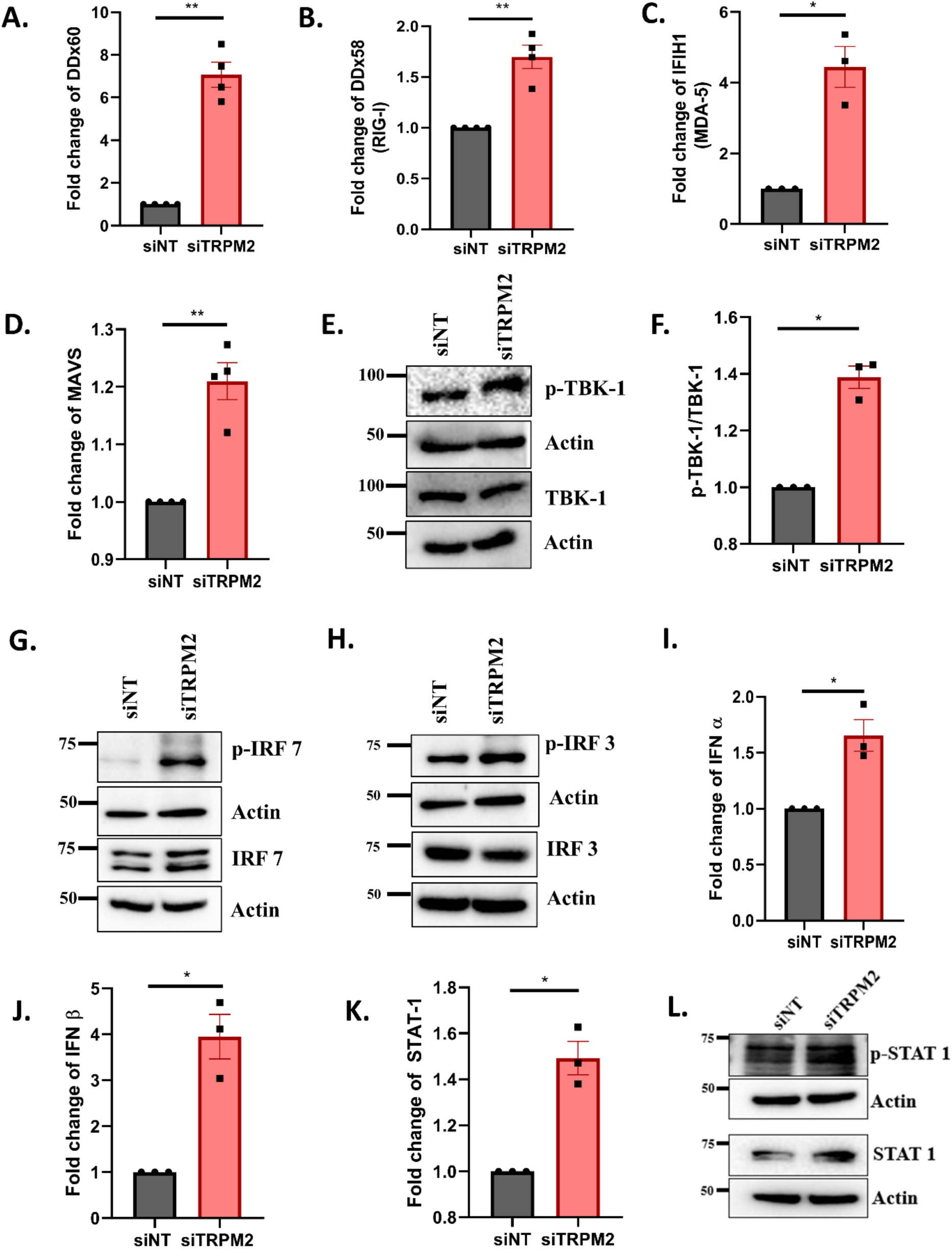
TRPM2 silencing triggers RLR signalling and generates type-1 IFN response leading to activate ISG15. (A) qRT-PCR analysis showing fold change of DDx60 mRNA levels in either siNT or siTRPM2 transfected LD day 6 cells (N=4). (B) qRT-PCR analysis showing fold change of RIG-I mRNA levels in either siNT or siTRPM2 transfected LD day 6 cells (N=4). (C) qRT-PCR analysis showing fold change of IFIH1(MDA-5) mRNA levels in either siNT or siTRPM2 transfected LD day 6 cells (N=3). (D) qRT-PCR analysis showing fold change of MAVS mRNA levels in either siNT or siTRPM2 transfected LD day 6 cells (N=4). (E) Representative western blot showing phospho-TBK-1(p-TBK-1) and TBK-1 protein levels in either siNT or siTRPM2 transfected LD day 6 cells (N=3). (F) Densitometric quantitation of ratio of phospho-TBK-1(p-TBK-1)/TBK-1 in either siNT or siTRPM2 transfected LD day 6 cells (N=3). (G) Representative western blot showing phospho-IRF-7 (p-IRF-7) and IRF-7 protein levels in either siNT or siTRPM2 transfected LD day 6 cells (N=3). (H) Representative western blot showing phospho-IRF-3(p-IRF-3) and IRF-3 protein levels in either siNT or siTRPM2 transfected LD day 6 cells (N=3). (I) qRT-PCR analysis showing fold change in the mRNA levels of IFN-α in either siNT or siTRPM2 transfected LD day 6 cells (N=3). (J) qRT-PCR analysis showing fold change in the mRNA levels of IFN-β in either siNT or siTRPM2 transfected LD day 6 cells (N=3). (K) qRT-PCR analysis showing fold change in the mRNA levels of STAT-1 in either siNT or siTRPM2 transfected LD day 6 cells (N=3). (L) Representative western blot showing phospho-STAT-1 and STAT-1 protein levels in either siNT or siTRPM2 transfected LD day 6 cells (N=3).

**Fig 8:**
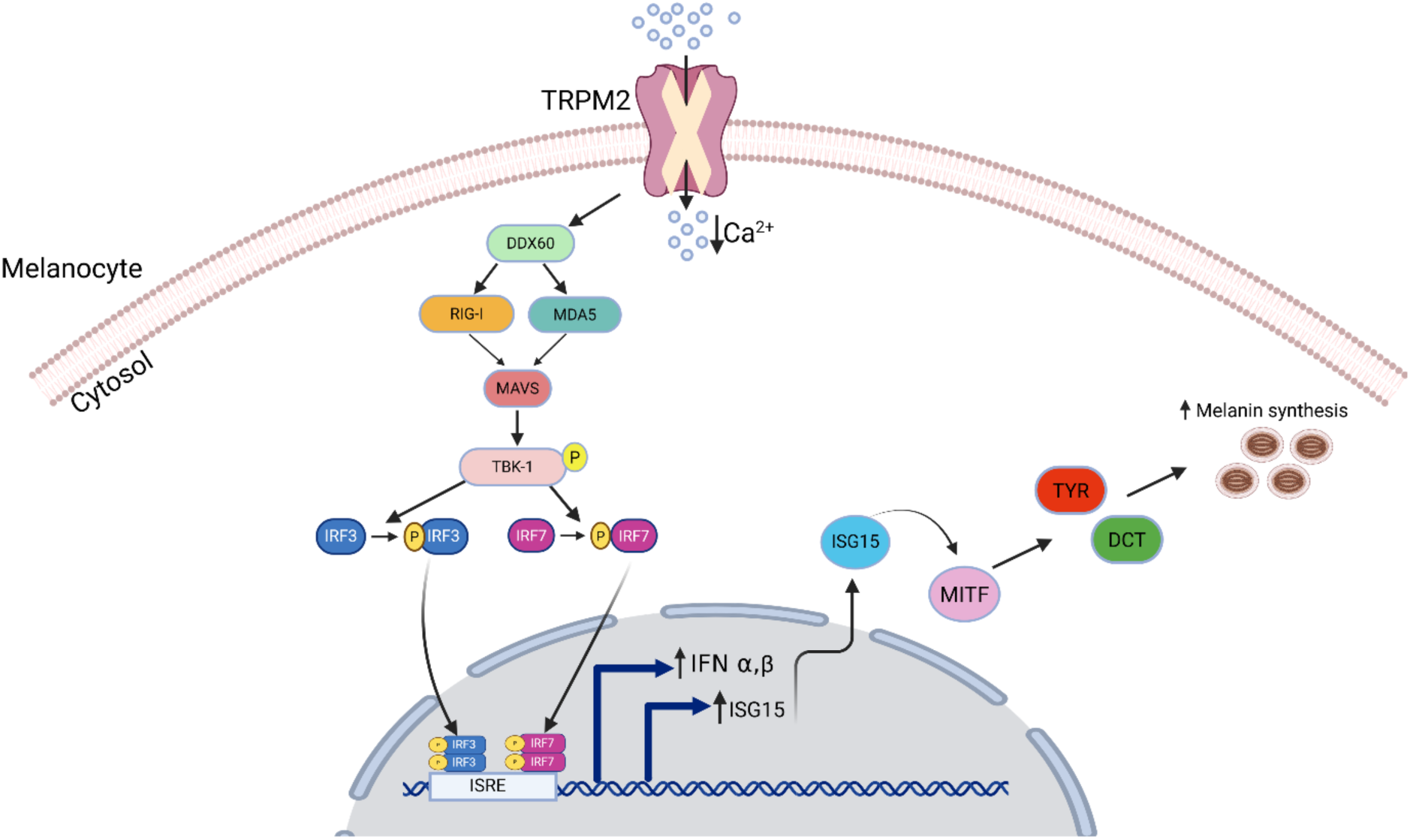
Graphical Summary. TRPM2 silencing triggers RLR signaling in melanocytes to generate a cell autonomous type-I-IFN response, which regulates melanogenesis within melanocytes. TRPM2 knockdown activates DDx60, a DExD/H box helicase, which acts as a sentinel for retinoic acid-inducible gene-I (RIG-I) like receptor (RLR) signaling. RIG-I signaling involves activation of RIG-I and MDA-5. Activated RIG-I and MDA-5 further activates downstream adaptor mitochondrial antiviral signaling protein (MAVS) and causes recruitment & activation of a kinase, TBK-1. TBK-1 in turn phosphorylates IRF-3 and IRF-7. Phosphorylated IRF3/7 induce type-I-IFNs alpha and beta for type-I-IFN pathway activation. Finally, type-I-IFN pathway activation induces ISG15, which regulates MITF expression positively and thereby enhances melanogenesis.

## Discussion

Skin Pigmentation plays a critical role in photoprotection and maintains skin homeostasis (Le et al., 2021; Lin & Fisher, 2007). While classical signaling pathways regulating the process of melanogenesis such as canonical MC1R dependent cAMP-PKA pathway, Wnt-β-catenin/MITF signaling and Ca^2+^ signaling have been extensively studied (Ahuja et al., 2025; Le et al., 2021; Sharma et al., 2020), emerging evidences also highlight the critical contribution of immunological molecules in melanogenesis (Fu et al., 2020; Natarajan, Ganju, Singh, et al., 2014). In line with this, several pigmentary disorders are increasingly recognized as manifestations of dysregulated immunity (Rashighi et al., 2014; Silpa-archa et al., 2017; Speeckaert et al., 2022; Tajik et al., 2024). Studies have highlighted the effect of immune-mediated cytotoxicity on melanocyte biology across the spectrum of pigmentary disorders such as lichen planus pigmentosus, halo nevus, alopecia areata (Fu et al., 2020; Rao et al., 2023; Silpa-archa et al., 2017; Speeckaert et al., 2022; Tajik et al., 2024) and vitiligo (H. Liu et al., 2024; Natarajan, Ganju, Singh, et al., 2014). Given the role of oxidative stress induced immune cascade activation, and melanocyte dysfunction in various pigmentary disorders, identification of molecular mediators that integrate pigmentary and immune pathways within melanocytes is essential. In this study, we identify TRPM2, a Ca^2+^ permeable channel as a novel regulator of pigmentation that links type-I-IFN signaling to pigmentation. Using a variety of *in-vitro* and *in vivo* experiments, our study elegantly demonstrates that TRPM2 negatively regulates pigmentation. Mechanistically, we reveal an immuno-pigmentary signaling module comprising TRPM2-type-I-IFN-ISG15-MITF axis that controls pigmentation. Given the strong association of immune dysfunction and pigmentary disorders, deciphering the role of TRPM2 in pigmentary conditions requires future investigation. Understanding the crosstalk of this axis with other canonical pathways that regulate MITF expression will help in developing therapeutic strategies against pigmentary disorders.

Ion channels have been increasingly recognized as integral regulators of melanocyte physiology wherein they influence intracellular/organellar Ca^2+^ dynamics, pH and Ca^2+^ homeostasis within melanosomal lumen, melanosome maturation and the cellular response to environmental cues (Jia et al., 2021; Oancea et al., 2009; Sharma et al., 2023; Wu et al., 2023). Nevertheless, studies have majorly focused on electrophysiological properties of ion channels and their involvement in regulating Ca^2+^ dynamics in context of melanogenesis with limited focus on their functional diversity such as their involvement in immune signaling (Devi et al., 2009; Jia et al., 2021; Oancea et al., 2009). Here, we extend this paradigm by demonstrating TRPM2 mediated type-I-IFN pathway activation in melanocytes positioning TRPM2 as critical regulator of immune dependent pigmentation process. One of the major strengths of this study is the robust validation of TRPM2 across multiple model systems. *In vitro* studies employing pharmacological inhibition, gene silencing and overexpression consistently demonstrate channel’s negative regulation over pigmentation (**Fig.1 and 2**). Notably, increase in pigmentation phenotype in primary melanocytes further strengthens this conclusion (**Fig. 2**). Importantly, *in vivo* studies using TRPM2 zebrafish morphants and global TRPM2 knockout mice (**Fig. 3**) are compelling underscoring the physiological relevance of TRPM2 in the context of human skin pigmentation.

Our unbiased RNA sequencing analysis identified ISG15 as a key and novel regulator of melanogenesis (**Fig. 4**). ISG15 is a member of the ubiquitin family (J. A. Kang et al., 2022; Mirzalieva et al., 2022) and has been extensively studied for its role in maintaining skin homeostasis, particularly in ulcerative skin lesions and skin inflammation (Buda et al., 2020; Y. Liu et al., 2023; Malik et al., 2022; Martin-Fernandez et al., 2020; Oyedepo et al., 2023). Several studies have implicated ISG15 deficiency or mutation in skin pathologies and inflammatory phenotype. A recent case wherein two siblings were reported with frequent episodes of skin ulcers and CNS calcifications, revealed a homozygous nonsense gene mutation in ISG15. *In vitro* characterization showed ISG15-deficient cells exhibiting hyper-inflammatory phenotype. Further deficiency of ISG15 in fibroblast cells resulted in increased ROS levels, and in keratinocytes showed retarded cell migration. Overall, this study showed the critical role of ISG15 in regulating the cell migration and maintaining tissue homeostasis, leading to predominant skin ulcers (Malik et al., 2022). In another case report, two pathogenic variants (c.285del and c.299_312del, NM_005101.4 GRCh37(hg19) of ISG15 were identified by whole exome sequencing. These variants resulted in complete ISG15 deficiency and manifesting into recurrent skin ulcers, cerebral calcifications and frequent episodes of pulmonary manifestations (Buda et al., 2020). Furthermore, 6 novel ISG15 mutations associated with skin lesions were recently identified in 5 patients resulting from elevated type -I-IFN response (Martin-Fernandez et al., 2020). Despite these finding explored and demonstrated the critical role of ISG15 in skin lesions, the involvement of ISG15 in regulating skin pigmentation has not been previously investigated. In this study, we demonstrate that TRPM2 silencing leads to enhanced ISG15 levels (**Fig. 5**). Importantly, ISG15 silencing reduces pigmentation whereas ISG15 OE enhances pigmentation. This collectively establish ISG15 as a novel positive regulator of melanogenesis (**Fig. 5**). Thus, this study determines a previously unrecognized role of ISG15 in pigmentation while providing a mechanistic link between interferon pathway and melanogenesis. Moreover, given the role of ISG15 in skin inflammation and skin lesions (Malik et al., 2022; Martin-Fernandez et al., 2020; Mirzalieva et al., 2022), this study provides a solid foundation to identify potential therapeutic avenue for skin diseases and interferonopathies (Y. Liu et al., 2023; Martin-Fernandez et al., 2020; Rao et al., 2023; Silpa-archa et al., 2017).

MITF has been widely recognized as the master regulator of melanogenesis where it regulates survival and differentiation of melanocytes (Nguyen & Fisher, 2019; Tanwar, Sharma, et al., 2022). MITF transcriptionally regulates a variety of genes crucial for melanocyte development and melanin synthesis (Nguyen & Fisher, 2019; Tanwar, Sharma, et al., 2022). In turn, MITF expression is modulated by several pathways and post translational modifications, which have been extensively characterized (Kawakami & Fisher, 2017). However, its modulation by an ISG/immune signaling remains largely unappreciated. Here, we identify that ISG15 regulates MITF expression in melanocytes (**Fig. 6**). The role of ISG15 in modulating MITF has not been described earlier. Therefore, to best of our knowledge, here we report for the first time, that ISG15 positively influences MITF expression. ISG15 silencing led to the reduction in MITF protein levels, while ISG15 OE elevated MITF protein levels, resulting in corresponding changes in pigmentation (**Fig. 6**). Consistent with MITF expression, ISG15 positively regulates TYR and DCT levels, both transcriptionally regulated by MITF (Nguyen & Fisher, 2019) (**Fig. 5**). Given that TYR and DCT are downstream targets of MITF in the melanin biosynthesis pathway, these observations demonstrate a direct axis wherein ISG15 modulates MITF and MITF in turn regulates TYR and DCT expression to control melanogenesis.

Interestingly, ISG15 regulates protein stability and turnover through ISGylation, which modulates protein degradation by competing with ubiquitination pathway (Desai et al., 2006; Fan et al., 2015; Okumura et al., 2008). ISG15 conjugation to its target proteins exhibits variability in their ubiquitylation status in a context-dependent manner. Several studies have reported that higher ISG15 signatures are linked to a reduction in ubiquitination level (Desai et al., 2006; Fan et al., 2015; Okumura et al., 2008). Additionally, the presence of mixed ISG15-Ub chains exert a negative influence on the degradation rate of ubiquitylated proteins (Fan et al., 2015). Conversely, emerging literature shows that ISG15 conjugation facilitates and promotes proteasomal degradation of its cellular targets via the proteasomal degradation pathway (Qu et al., 2023). Here, our study demonstrates that ISG15 antagonizes global ubiquitination in melanocytes (**Fig. 6**). ISG15 modulation via silencing and overexpression led to a respective increase and decrease in the ubiquitination status of B16 cells. In similar assays, we found that ISG15 can positively regulate MITF levels by limiting its degradation (**Fig. 6**). In future studies, it would be interesting to identify the specific sites of ubiquitination on MITF and to examine the potential E3 ubiquitin ligases or deubiquitinases that work downstream of ISG15 in regulating MITF levels.

Literature suggests a role of oxidative stress-type-II IFN axis in melanogenesis (H. Liu et al., 2024; Rashighi et al., 2014) but the significance of type-I-IFN in pigmentation biology remains largely unexplored. Our study demonstrates an ion channel mediated autonomous activation of the RLR signaling and type-I-IFN response within melanocytes. Our unbiased transcriptomics data reveals that TRPM2 silencing activates a DExD/H box helicase 60, leading to RLR signaling activation and subsequent phosphorylation of TBK-1, IRF3 and IRF7 (**Fig. 7**). This culminates in sustained type-I-IFN induction, predominantly IFN-β causing ISG15 upregulation (**Fig. 7**). Additionally, TRPM2 silencing mediated type-I-IFN induction in melanocytes is particularly notable as both TRPM2 and type-I-IFN have been widely associated with immunity and inflammation but not in pigmentation (Beceiro et al., 2017; Knowles et al., 2011; McNab et al., 2015; Syed Mortadza et al., 2015). In this study, we revealed that TRPM2 silencing leads to type-I-IFN activation and downstream ISG15 induction to govern pigmentation.

The pathways activating type-I-IFN response majorly converge on a set of pattern recognition receptor (PRR) system that is responsible in detecting the exogenous pathogen associated molecular patterns (PAMPs) and endogenous damage associated molecular patterns (DAMPs). These are further categorized into families such as toll like receptors (TLRs), retinoic acid-inducible gene-I (RIG-I) like receptors (RLR), nucleotide oligomerization domain (NOD)-like receptors (NLR) and cyclic GMP-AMP synthase(cGAS)-stimulator of IFN genes (STING) (Li & Wu, 2021). In this study, we investigated the potential activation of cGAS-STING pathway in addition to RLR signaling downstream of TRPM2 silencing as these pathways have recently emerged as critical regulators of innate immunity and inflammation (Mathavarajah et al., 2019). Further, a recent study demonstrated a role of TRPM2 in osteoarthritis (OA) wherein TRPM2 mediated feed forward loop involving Ca^2+^ cGAS-STING-NF-kB axis was reported (Sun et al., 2025). Activation of cGAS-STING pathway requires sensing of pathogen derived or endogenous double stranded DNA (dsDNA) released from nucleus or mitochondrial (mt) into the cytosol during cellular stress, cell damage or injury (Kim et al., 2023). Thus, we assessed mtDNA release following TRPM2 silencing and detected no cytosolic mtDNA (**Supplementary Fig. 3E**) indicating that this pathway is not activated. Consistently, cGAS and STING protein expression analysis and phosphorylation revealed no significant changes (**Supplementary Fig. 3F-H**) confirming that cGAS-STING axis remains inactive downstream of TRPM2 silencing. In future, it would be worth investigating the role of other pathways in melanogenesis.

Our finding of TRPM2-type-I-IFN-ISG15-MITF axis is significant as literature suggests that MITF expression regulates melanoma progression (Kawakami & Fisher, 2017). Further, ISG15 was recently reported to be constitutively produced by melanoma cells and to promote melanoma progression (Filderman et al., 2024; Padovan et al., 2002). Interestingly, loss of TRPM1 expression is also associated with melanoma progression wherein MITF directly regulates TRPM1 transcription (Kawakami & Fisher, 2017; Miller et al., 2004). Further, a study reported down-regulation of full-length TRPM2 and upregulation of anti-sense transcripts in melanoma cell line (Orfanelli et al., 2008). While TRPM1 and MITF have been extensively studied in melanoma biology, the regulation of melanoma via TRPM2 and MITF crosstalk remains poorly understood. Hence, in future, it would be worth to study the TRPM2-ISG15-MITF axis in melanoma and other cancers.

Taken together, our study reveals a critical role of TRPM2 in regulating pigmentation. Our in *vitro* studies with TRPM2 inhibition, gene silencing and overexpression in B16 LD model and in primary human melanocytes elegantly demonstrate that TRPM2 negative regulates pigmentation. Further *in vivo* studies with TRPM2 zebrafish morphants and TRPM2^-/-^ mice model validate *in vitro* findings. Next, through unbiased RNA-sequencing, we uncovered ISG15 as a crucial positive regulator of pigmentation working downstream of TRPM2. Mechanistically, we here reveal a novel TRPM2-type-I-IFN-ISG15-MITF axis that orchestrates pigmentation. Since TRPM2, ISG15, and MITF play a very important role in inflammation, immunity and melanoma; the signaling module reported in this study may have broader implications beyond pigmentation, including immune-disorders, melanoma and immunotherapy.

## Supplementary Figures

**Supplementary Fig 1 Supporting main Fig 2:**
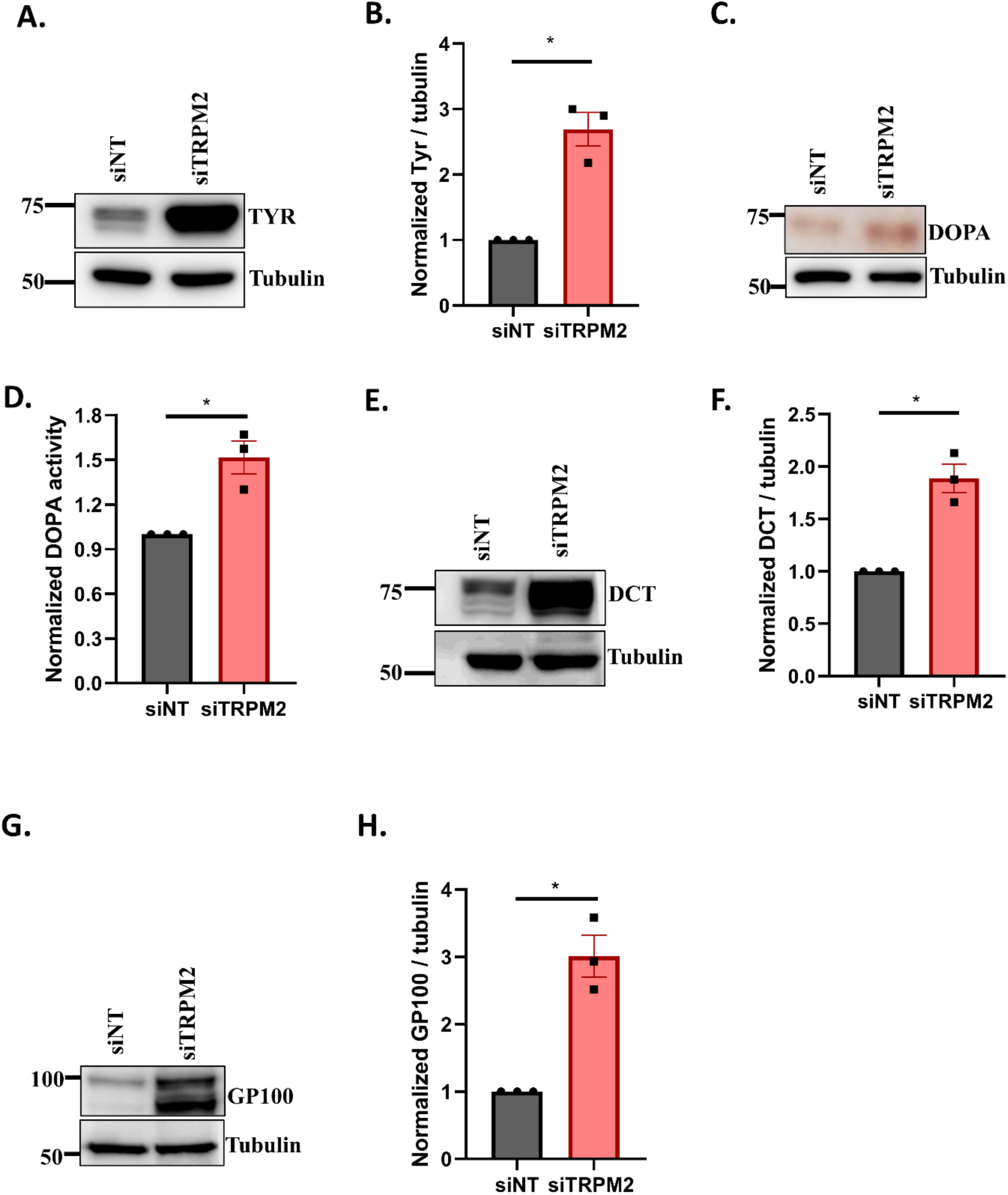
TRPM2 loss & gain of function studies establish it as a negative regulator of pigmentation. (A) Representative western blot of TYR from either siNT or siTRPM2 transfected LD day 6 cells (N=3). (B) Densitometric quantitation of TYR protein levels from either siNT or siTRPM2 transfected LD day 6 cells (N=3). (C) Representative in-gel image of DOPA assay to measure TYR activity from either siNT or siTRPM2 transfected LD day 6 cells (N=3). (D) Densitometric quantitation of DOPA assay from either siNT or siTRPM2 transfected LD day 6 cells(N=3). (E) Representative western blot of DCT from either siNT or siTRPM2 transfected LD day 6 cells (N=3). (F) Densitometric quantitation of DCT protein levels from either siNT or siTRPM2 transfected LD day 6 cells (N=3). (G) Representative western blot of GP100 from either siNT or siTRPM2 transfected LD day 6 cells (N=3). (H) Densitometric quantitation of GP100 protein levels from either siNT or siTRPM2 transfected LD day 6 cells (N=3).

**Supplementary Fig 2 Supporting main Fig 2:**
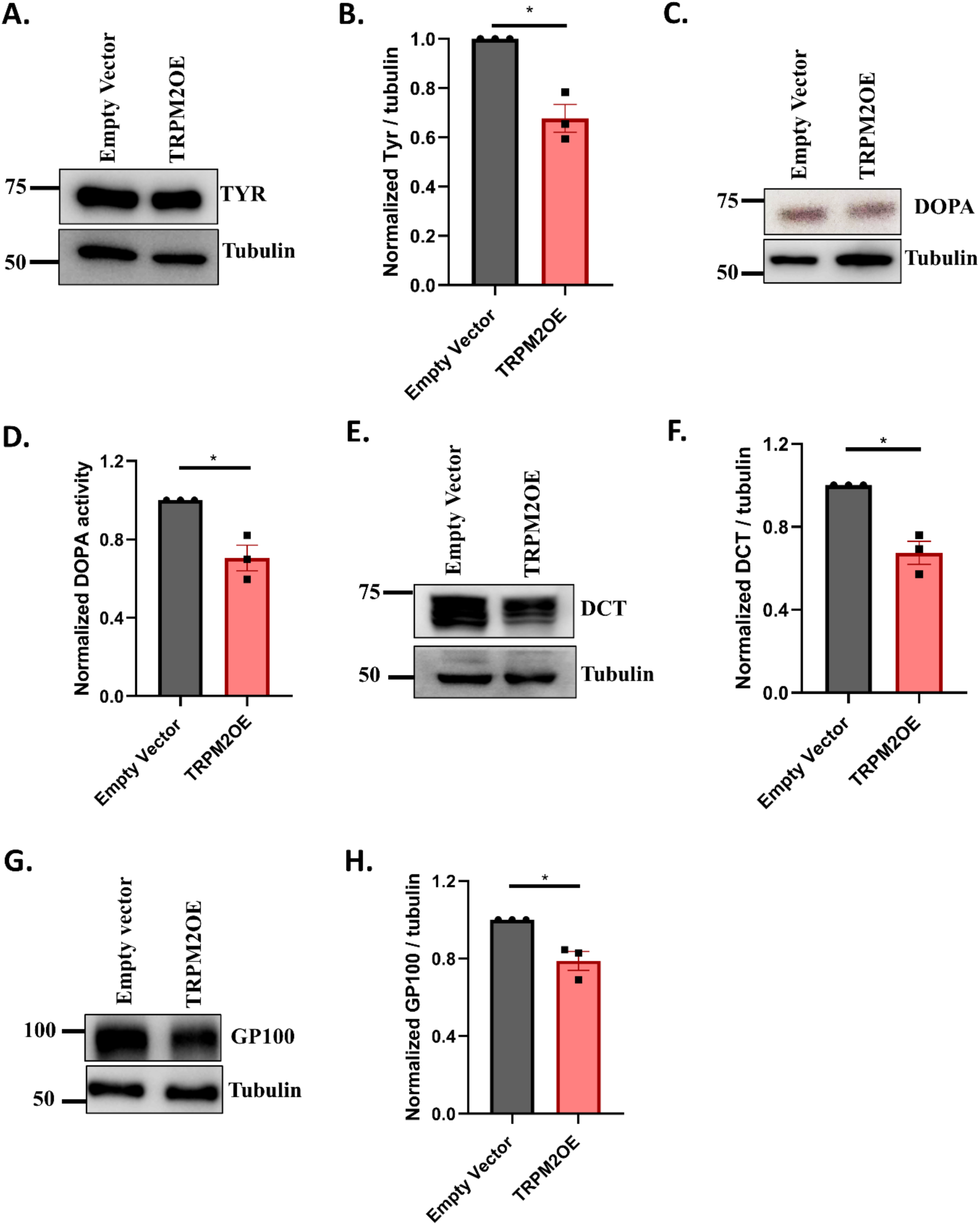
TRPM2 loss & gain of function studies establish it as a negative regulator of pigmentation. (A) Representative western blot of TYR from either EV or TRPM2 transfected LD day 6 cells (N=3). (B) Densitometric quantitation of TYR protein levels from either EV or TRPM2 transfected LD day 6 cells (N=3). (C) Representative in-gel image of DOPA assay to measure TYR activity from either EV or TRPM2 transfected LD day 6 cells (N=3). (D) Densitometric quantitation of DOPA assay from either EV or TRPM2 transfected LD day 6 cells (N=3). (E) Representative western blot of DCT from either EV or TRPM2 transfected LD day 6 cells (N=3). (F) Densitometric quantitation of DCT protein levels from either EV or TRPM2 transfected LD day 6 cells (N=3). (G) Representative western blot of GP100 from either EV or TRPM2 transfected LD day 6 cells (N=3). (H) Densitometric quantitation of GP100 protein levels from either EV or TRPM2 transfected LD day 6 cells (N=3).

**Supplementary Fig 3 Supporting main Fig 7:**
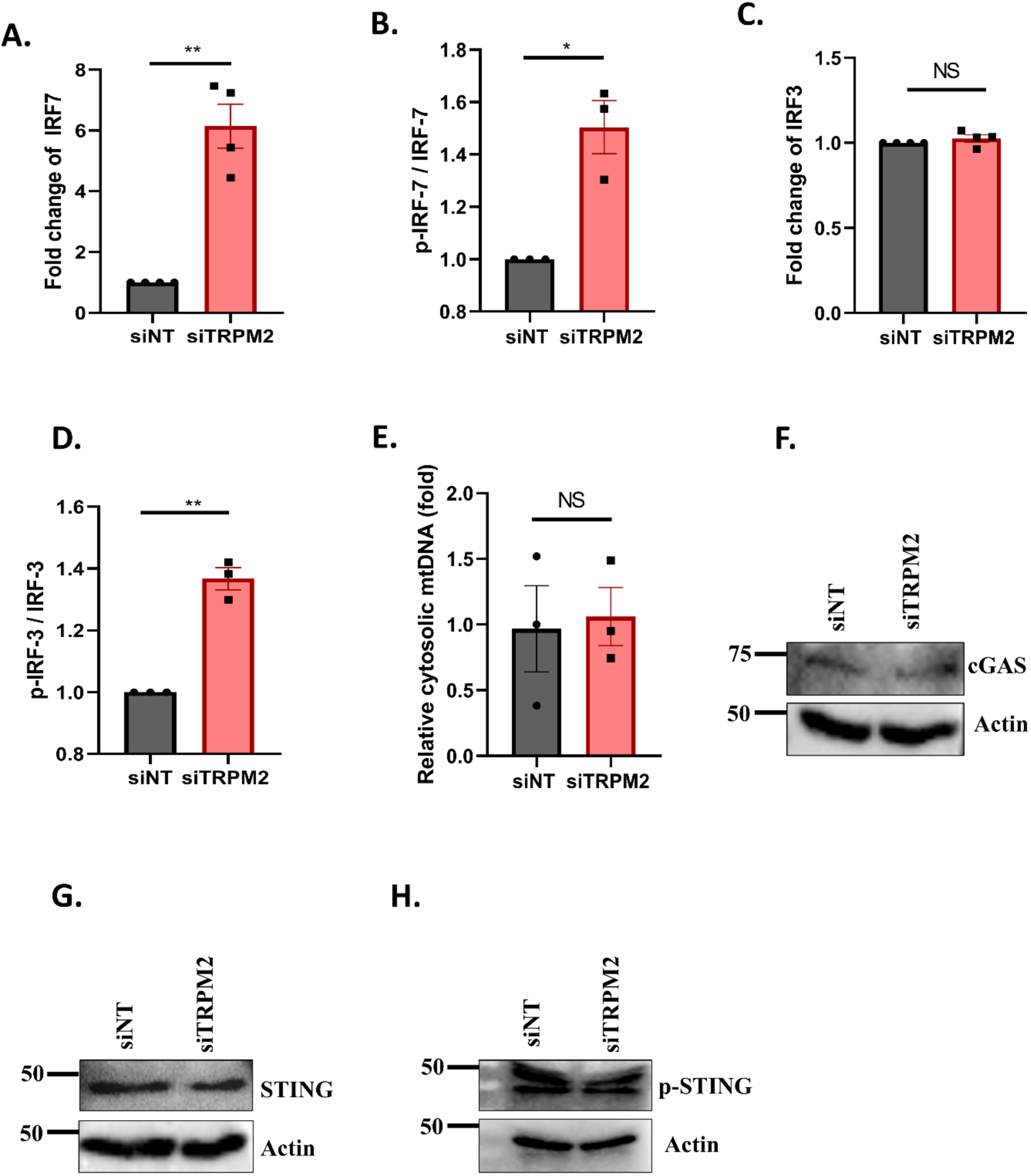
TRPM2 silencing triggers RLR signaling to generate type-1 IFN response leading to the ISG15 activation. (A) qRT-PCR analysis showing fold change in IRF-7 mRNA levels in either siNT or siTRPM2 transfected LD day 6 cells (N=4). (B) Densitometric quantitation of ratio of phospho-IRF-7(p-IRF-7)/IRF-7 in either siNT or siTRPM2 transfected LD day 6 cells (N=3). (C) qRT-PCR analysis showing fold change in IRF-3 mRNA levels in either siNT or siTRPM2 transfected LD day 6 cells (N=4). (D) Densitometric quantitation of ratio of phospho-IRF-3/(p-IRF-3) in either siNT or siTRPM2 transfected LD day 6 cells (N=3). (E) qRT-PCR analysis showing fold change of cytosolic mtDNA detected in either siNT or siTRPM2 transfected LD day 6 cells (N=3) (F) Representative western blot showing cGAS protein levels in either siNT or siTRPM2 transfected LD day 6 cells (N=3). (G) Representative western blot showing STING protein levels in either siNT or siTRPM2 transfected LD day 6 cells (N=3). (H) Representative western blot showing phospho-STING protein levels in either siNT or siTRPM2 transfected LD day 6 cells (N=3).

## Materials and Methods

### Cell culture

B16-F10 (from ATCC) were cultured under standard conditions (37°C and 5% CO2) in DMEM (Sigma, Bangalore, India) supplemented with 10% Fetal Bovine Serum (FBS). Cell culture reagents such as Trypsin, Dulbecco’s Phosphate Buffer Saline (DPBS), Versene, Fetal Bovine Serum (FBS) were obtained from Invitrogen, Waltham, MA, USA. Lightly pigmented (LP) primary human melanocytes were procured from Invitrogen, Waltham, Massachusetts, USA. Cells were grown in Medium 254 supplemented with human melanocyte growth supplement-2 and maintained at 37°C in a humidified incubator with 5% CO_2_ atmosphere. Cell maintenance and subculturing were carried out as per the manufacturer’s recommendations.

### B16 Low-Density (LD) pigmentation-oscillator model

Non-pigmented B16-F10 cells were seeded at the density of 100 cells/cm2 in DMEM supplemented with 10% Fetal Bovine Serum as described earlier (Motiani et al., 2018; Natarajan, Ganju, Ramkumar, et al., 2014) and were incubated at 37°C in a 5% CO2 incubator for 6-7 days wherein these cells showed an autonomous pigmentation.

### Reagents and plasmids

N-(p-amyl cinnamoyl) anthranilic acid (ACA)-CAS 110683-10-8 was procured from Sigma. ACA was dissolved in dimethyl sulfoxide (DMSO). 20uM working concentration of ACA was used in the cell culture experiments. The antibodies used were procured from Abcam, Cell Signaling Technology (CST) and Santa Cruz (SC) Biotechnology. *N*-Phenylthiourea (PTU) was procured from Sigma. L-DOPA substrate was procured from Sigma. Plasmids used in this study were procured from Addgene and are listed in the Table 1.

| Plasmid | Catalog Number |
| --- | --- |
| TRPM2 overexpression plasmid | #53920 |
| ISG15 overexpression plasmid | #80404 |

### siRNA-based Transfections

B16-F10 mouse melanoma cells were seeded at the density of 100 cells/cm2 in DMEM-HG growth medium with added antibiotics in T75 flasks, and on day 3, siRNA transfections were performed using Dharmafect in the ratio of 1:3. The transfections were done in OptiMEM for efficient transfection and post 4-6 hours of transfection, the transfection medium was replaced with LD day 3 medium. Flasks were incubated at 37°C in a 5% CO2 incubator till day 6 termination. At day 6, cells were harvested, and downstream processing was performed. The siRNAs were procured from Dharmacon.

The catalog numbers of siRNAs used in the study are included in the table 2

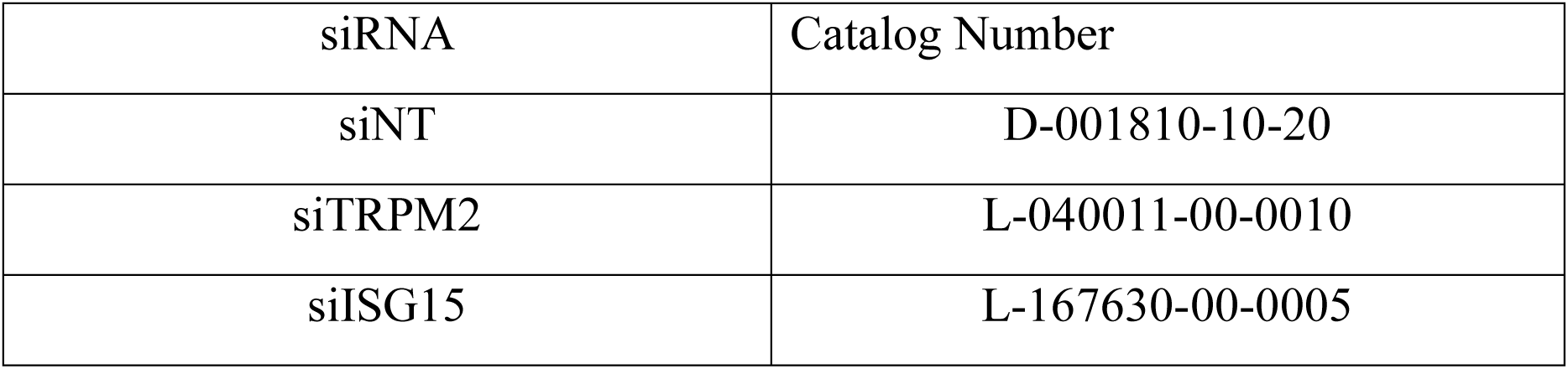

| siRNA | Catalog Number |
| --- | --- |
| siNT | D-001810-10-20 |
| siTRPM2 | L-040011-00-0010 |
| siISG15 | L-167630-00-0005 |

### Western blotting

B16-F10 cell protein lysate was prepared with NP-40 lysis buffer and were supplemented with protease inhibitors before keeping for overnight incubation at -80 degrees for lysis. Following lysis, protein concentration was determined and approximately 50-100 ug of protein was loaded onto gels for western blot analysis. The separated proteins were then transferred onto PVDF membranes via electroblotting. Membranes were blocked in 5% non-fat dry milk prepared in 1X TBST to prevent non-specific binding. Subsequently, blots were kept for overnight incubation at 4°C with specific primary antibodies. Primary antibodies were mainly obtained from Abcam and Cell Signaling Technology (CST) and Santa Cruz (SC). Following primary incubation, membranes were washed and then incubated for 2 hours at room temperature with horseradish peroxidase (HRP)-conjugated secondary antibodies. After incubation with secondary antibodies, membranes were washed, and the signal of bound antibody was detected using the enhanced chemiluminescence (ECL) kit. Densitometric analysis of blots was done using ImageJ software, and data are graphically represented as mean ± SEM.

The catalog number and company name for the antibodies is provided in Table 3

| **Antibody** | **Company** | **Catalog Number** | **Species** |
| --- | --- | --- | --- |
| TRPM2 | Abcam | ab11168 | Mouse |
| ISG15 | Santa Cruz | sc-166755 | Mouse |
| Ubiquitin | Santa Cruz | sc-271289 | Mouse |
| MITF | Santa Cruz | sc-56725 | Mouse |
| DCT | Abcam | ab74073 | Mouse |
| TYR | Abcam | ab170905 | Mouse |
| GP100 | Abcam | ab137078 | Mouse |
| Tubulin | Abcam | ab21058 | Mouse |
| Actin | Sigma-Aldrich | A3854 | Mouse |
| TBK-1 | CST | 38066 | Mouse |
| p-TBK-1 | CST | 5483 | Mouse |
| IRF-7 | CST | 72073 | Mouse |
| p-IRF-7 | CST | 24129 | Mouse |
| IRF-3 | CST | 4302 | Mouse |
| p-IRF-3 | CST | 29047 | Mouse |
| cGAS | CST | 79978 | Mouse |
| STING | CST | 13647 | Mouse |
| p-STING | CST | 72971 | Mouse |

### Melanin Content assay

Melanin content assay was performed as described earlier (Sultan et al., 2022; Tanwar, Saurav, et al., 2022). In summary, cells were lysed in 1 N NaOH by heating at 80°C for 2 h, and then, absorbance was measured at 405 nm. The melanin content was determined by interpolating the sample absorbance readings on a standard curve (µg/ml) generated with synthetic melanin.

### DOPA assay

Tyrosinase enzyme activity in cell lysates was assessed by performing the DOPA assay as previously described by Fuller et al. (2000). In brief, cell lysates were prepared in NP-40 lysis buffer, and an equal amount of protein for different conditions was run on a gel under non-reducing/native conditions. The gel was then incubated in a phosphate buffer containing the tyrosinase substrate L-DOPA (Sigma Chemicals, Bangalore, India). Enzyme activity corresponded to the formation of a black color pigment.

### Overexpression with Plasmids

B16-F10 mouse melanoma cells were seeded at the density of 100 cells/cm2 in DMEM-HG growth medium with added antibiotics in T75 flasks, and on day 3, plasmid transfections were performed using lipofectamine reagent in the ratio of 1:2. The transfections were done in OptiMEM for efficient transfection and post 4-6 hours of transfection, the transfection medium was replaced with LD day 3 medium. Flasks were incubated at 37°C in a 5% CO2 incubator till day 6 termination. On day 6, cells were harvested, and downstream processing was performed. Human TRPM2 OE plasmid (#53920) and human ISG15 OE plasmid (#80404) were obtained from Addgene.

### RNA sequencing

Total RNA was extracted from siNT, siTRPM2, pCDNA3.1 and TRPM2OE conditions in LD day 6 B16 cells using the Qiagen RNeasy kit following the manufacturer’s instructions. RNA was sent out to Clevergene, Bengaluru, India. Library preparation and high-throughput sequencing were performed using Illumina sequencers to generate paired-end results. Quality-based gene filtering was done with FastQC and MultiQC software, and quality-filtered reads were aligned with the reference genome GRCm39. Firstly, Gene expression values were obtained as read counts and normalized counts were generated. Fold change was calculated for siTRPM2 using non-targeting siNT as control. Log2 fold change value of more than 1 was used to filter the significantly upregulated genes, and genes with log2 fold change value less than - 1 were counted as significantly downregulated in TRPM2 silenced samples.

### Detection of cytosolic mtDNA via qPCR

Release and detection assay for mtDNA into the cytosol was performed following the previously established methodology(Bryant et al., 2022). Briefly, B16 cells were transfected with siNT and siTRPM2 in LD model at day 3 and were harvested at day 6 of LD. Crude cytosolic, mitochondrial and nuclear fractions were obtained as per the manufacturer instructions. Total DNA and proteins were extracted per fractions from both siNT and siTRPM2 transfected cells. Subsequently, purity of isolated fractions was confirmed with SDS-PAGE using actin as cytosolic marker, cytochrome c oxidase subunit (COX4, mouse monoclonal antibody, #11967, 1:1000) as mitochondrial marker and Lamin A/C (rabbit monoclonal antibody, #4L8Q0, 1:1000) as nuclear marker. Further, qPCR was performed to determine the release of mtDNA into the cytosol using TERT as nuclear marker, D-LOOP as mitochondrial marker. The primer sequences used are listed in the table.

### Zebrafish husbandry

Zebrafish used in this study were housed in the Laboratory of Calciomics and Systemic Pathophysiology at the Regional Centre for Biotechnology, India, with proper standard ethical protocols approved by the Institutional Animal Ethics Committee of Regional Centre for Biotechnology, India, with care to minimize animal suffering.

### Morpholino design and microinjections

Antisense morpholino (MO) oligonucleotides targeting the TRPM2 gene of zebrafish were designed to inhibit translation and procured from Gene Tools (USA). The MOs were reconstituted in nuclease-free water (Ambion, USA) following the manufacturer’s protocol to obtain a 1 mM stock solution, which was stored at −20°C until further use. For the experiments, zebrafish embryos were treated with *N*-Phenylthiourea (PTU) for 24 hours post 1 day of fertilization to prevent pigmentation. MO injections were then performed at 48 hours post-fertilization (hpf), after which the embryos were transferred to normal E3 medium and maintained at 28°C to allow pigmentation recovery. 50uM MO concentration was used for experiments. A scrambled MO served as a negative control. Embryos were examined for pigmentation phenotypes at 72 hpf, and images were captured using a Nikon microscope.

The morpholino sequences used in this study are listed in Table 4.

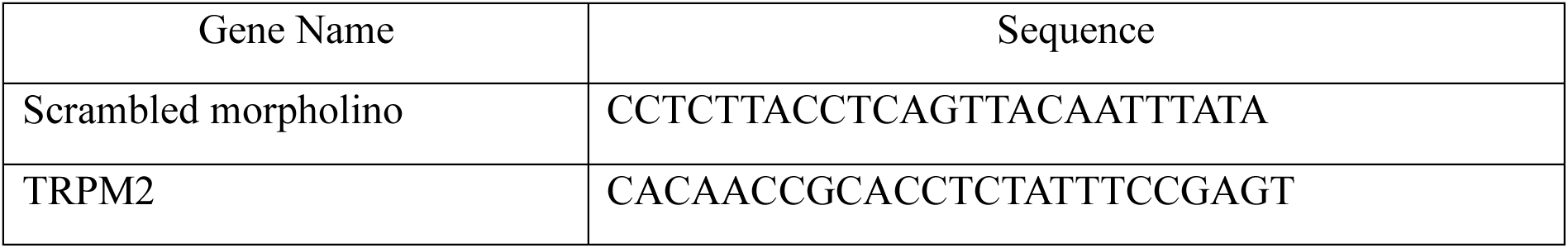

### qRT-PCR analysis

Total RNA was extracted from siNT, siTRPM2 conditions of LD day 6 B16 cells using the Qiagen RNeasy kit following the manufacturer’s instructions. 1ug total RNA sample was used to synthesize cDNA using the high-capacity cDNA reverse transcription kit from ThermoFisher (Waltham, MA, USA, Catalog#4368814). Real-time PCR (qRT-PCR) was performed using a 10ul reaction volume in 96 well plates by using SYBR green (Catalog #RR420A, Takara) according to the manufacturer’s protocol in Quant Studio 6 Flex from Applied Biosystems. The data was analyzed by normalizing the expression of siTRPM2, siNT cells with the housekeeping gene GAPDH using Quant Studio real-time PCR software version 1.3. Gene-specific primers were obtained from Eurofins.

The sequences of primers used in the study are listed in the Table 5.

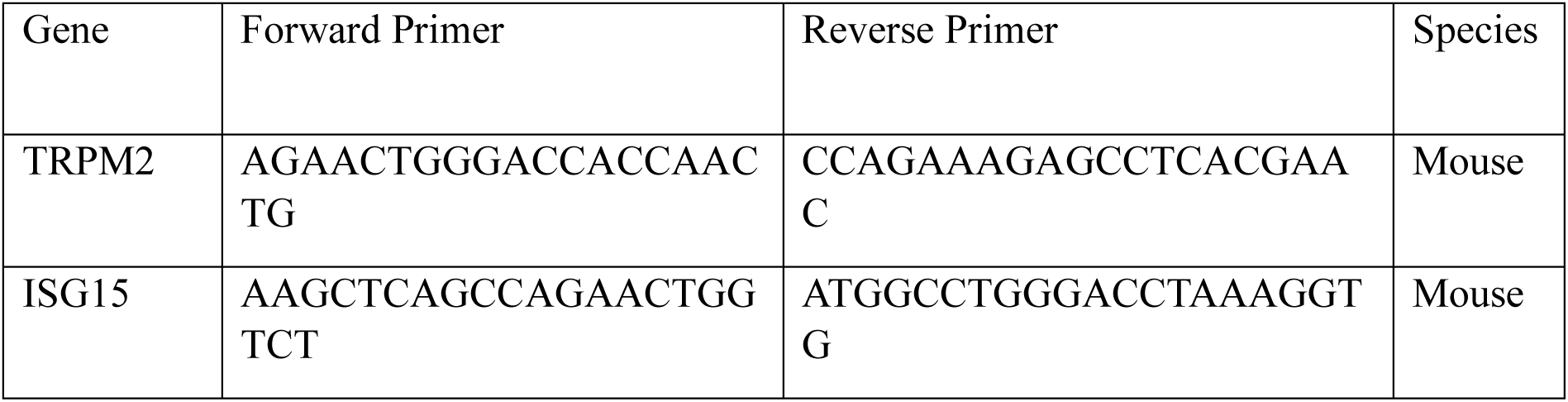
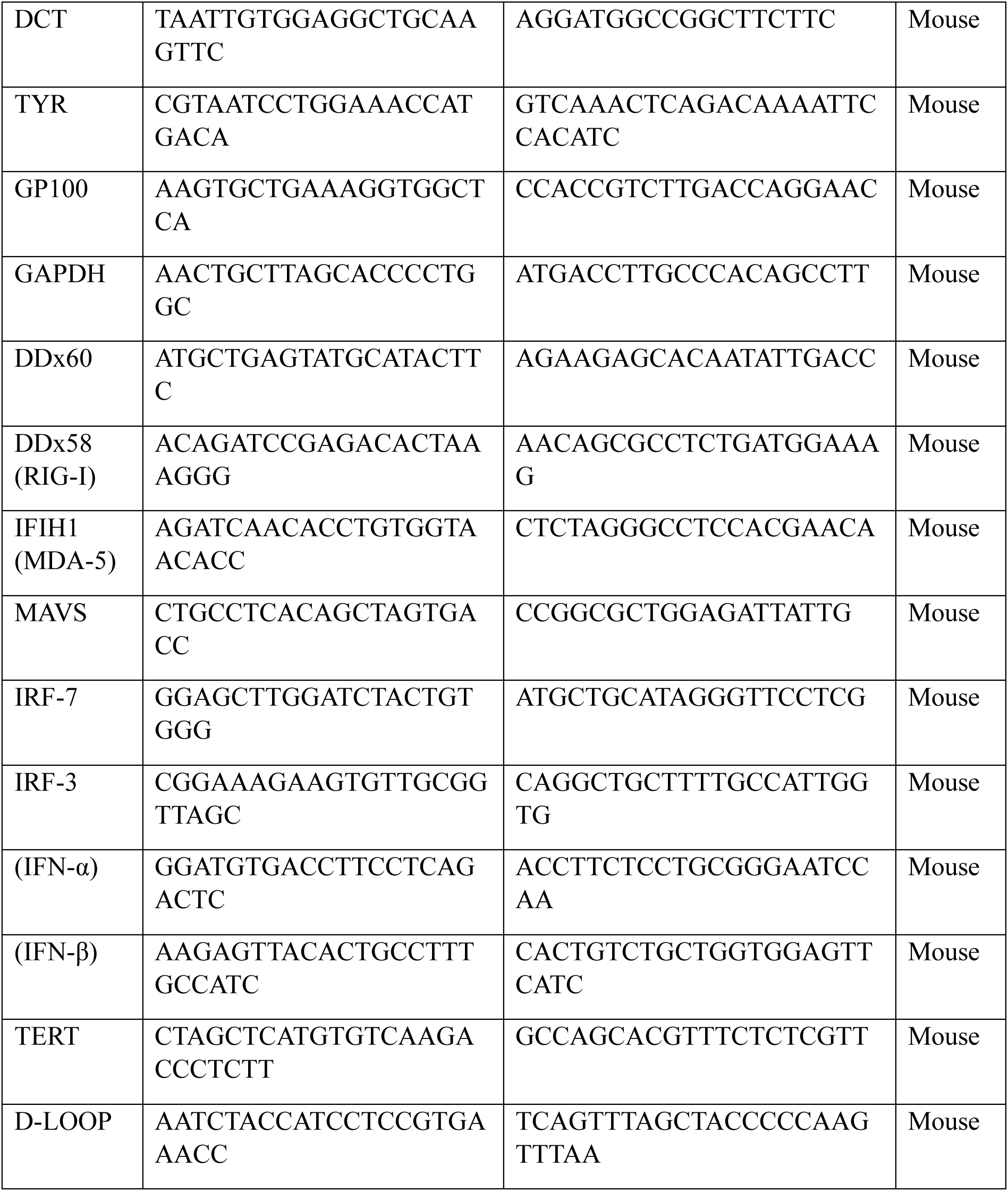

### Statistical Analysis

All statistical analysis was performed using GraphPad Prism 8 software. All the experiments were performed at least three times. Data are presented as mean ± SEM, and one sample t-test was performed to determine the statistical significance. p-value < 0.05 was considered as significant and is presented as “*”; p-value < 0.01 is presented as “**”and p-value < 0.001 is presented as “***”.

## Acknowledgements

This work was supported by the DBT/Wellcome Trust India Alliance Fellowship (IA/I/19/2/504651) to RKM. RKM also acknowledges funding support from RCB Institutional Core funding (R25226). IA acknowledges funding support from NIDCR-DIR, National Institutes of Health (NIH), United States (Z01-DE00438-38). The authors thank Dr. Manjula Kalia (Regional Centre for Biotechnology, India) for sharing reagents and antibodies. The authors also thank members of the Motiani laboratory for the critical discussions. NS acknowledges her Junior and Senior Research Fellowship from DBT, India. AT acknowledges his Junior Research Fellowship from DBT, India.

## Authors Contribution

Nutan Sharma: Methodology, Investigation, Visualization, Formal analysis, Writing-Original draft preparation. Abhishek Tanwar: Investigation, Visualization. Changyu Zheng: Investigation, Visualization. Indu Ambudkar: Visualization, Supervision. Rajender K Motiani: Conceptualization, Supervision, Writing-Original draft preparation, Reviewing and Editing, Project administration, Funding acquisition.

